# Machine Learning Analysis of the Human Initiator Region Reveals Key Features of Different Types of Core Promoters

**DOI:** 10.1101/2025.11.21.689830

**Authors:** Torrey E. Rhyne-Carrigg, Long Vo ngoc, Claudia Medrano, Kassidy E. Gillespie, James T. Kadonaga

## Abstract

The initiator (Inr) is the starting point for the transcription of many genes. Here, we generated highly predictive machine learning models of the human Inr region, and determined that the Inr is present in about 60% of natural promoters, identified a novel TATA-specific Inr, and detected the overlapping but functionally distinct TCT motif. Quantitative genome-wide analyses revealed a strict and synergistic interaction between the Inr and DPR, a duality between the TATA and DPR, a flexible and sometimes independent function of the TATA box in relation to the Inr, and different properties of the TCT motif in humans and *Drosophila*.

## Introduction

The RNA polymerase II transcription system is responsible for the expression of protein-coding genes as well as many noncoding RNAs in eukaryotes. The initiation of RNA polymerase II transcription is mediated by the core promoter, which is the stretch of DNA that is approximately −40 to +40 nucleotides relative to the +1 transcription start site (TSS) (for reviews, see Smale and Kadonaga 2003; Vo ngoc et al. 2017a; Haberle et al. 2018; Roeder 2019; Schier and Taatjes 2020; Sloutskin et al. 2021; Archuleta et al. 2024). The core promoter, the “gateway to transcription”, is at a key strategic position in the transcription process, as it is the site of convergence of the signals that lead to initiation.

All core promoters do not function by the same mechanism, but rather, different core promoters can have different activities that are determined by the presence or absence of DNA sequence elements such as the TATA box, downstream promoter region (DPR), initiator (Inr), and TCT motif (Supplemental Fig. S1). [The DPR is a unified version of the MTE and DPE elements (Vo ngoc et al. 2020). We will use the term DPR when referring to the DPE or DPR.] In human promoters, the estimated abundances of the TATA, Inr, and DPR are 15-23%, 40-56%, and 25-34%, respectively (Vo ngoc et al. 2017b, 2020). There are no universal core promoter elements, and many human promoters lack any of the known elements. Importantly, the individual core promoter elements confer specific transcriptional regulatory properties. For instance, in *Drosophila*, TATA-specific and DPR-specific enhancers have been identified (Butler and Kadonaga 2001; Juven-Gershon et al. 2008).

The Inr is the central focal point for the initiation of transcription of many genes. In an early analysis of eukaryotic promoters, a YYCA_+1_YYYYY consensus was identified at the TSS (Corden et al. 1980). This motif was incisively demonstrated to be a core promoter element and termed the Inr by Smale and Baltimore (1989). The Inr is recognized by the TFIID complex (Kaufman and Smale 1994; Purnell et al. 1994; Verrijzer et al. 1995), which also binds to the TATA box and DPR. Because of its abundance and correspondence with the +1 TSS, it is essential to have an accurate and reliable predictive model of the Inr.

In our previous studies of the core promoter, we used a combination of highthroughput analysis of randomized promoter elements (HARPE) and machine learning (SVR, support vector regression) to generate SVR models of the human TATA box and the human and *Drosophila* DPR (Vo ngoc et al. 2020, 2023). These SVR models provide objective, quantitative, data-based predictions of the activity of core promoter motifs and have been found to be more effective at predicting transcriptional activity than consensus sequence-based methods (Vo ngoc et al. 2020). One key advantage of the HARPE-SVR approach is that it involves the analysis of several hundred thousand different DNA sequence variants, which provide much more information than sequences from ∼10,000-30,000 natural promoters. In addition, in each HARPE assay, we vary the DNA sequences in only the target region of interest in an otherwise constant promoter backbone. This approach provides high quality information on a specific segment of the promoter. Thus, in this work, we focused on the Inr, which is the key core promoter element between the TATA and DPR, and generated multiple different SVR models of the human Inr region in different promoter and transcriptional (*i.e.*, biochemical vs. cell-based) contexts. The Inr SVR models, in conjunction with SVR models of the TATA and DPR, revealed key features of different types of core promoters that contain the Inr or the overlapping but functionally distinct TCT motif.

## Results and Discussion

### SVR models of the human Inr

To generate predictive models of the human Inr, we performed HARPE (Fig. 1A) and SVR analyses by the method of Vo ngoc et al. (2020). To this end, we first measured the biochemical transcription activity of ∼500,000 Inr sequence variants in which a 10-nt region (from −4 to +6 relative to the +1 TSS in the SCP1 promoter backbone) was randomized in a DPR-driven promoter as well as in a separate TATA-driven promoter (Fig. 1A). Because DPR- and TATA-dependent transcription occur by distinct mechanisms (see, for example: Willy et al. 2000; Hsu et al. 2008; Vo ngoc et al. 2020), we analyzed the properties of the Inr in the two different contexts. The HARPE data from independent replicates were found to be reproducible (Supplemental Table 1). A small subset of the sequence variants exhibited high transcriptional activity (Fig. 1B), and HOMER (Heinz et al. 2010) analysis of the top 0.1% of variants revealed a motif that strongly and somewhat surprisingly (in relation to previous human Inr consensus sequences; see, for example, Vo ngoc et al. 2017b) resembles the *Drosophila* Inr consensus, TCA_+1_KTY or TCA_+1_GTY (Fig. 1C, Supplemental Fig. S2; see, for example: Hultmark et al. 1986; Ohler et al. 2002, Fitz-Gerald et al. 2006). It is important to note, however, that TCA_+1_KTY is the consensus of the most active Inr sequences and should not be interpreted to be representative of typical Inr elements in natural promoters.

**Figure 1.**
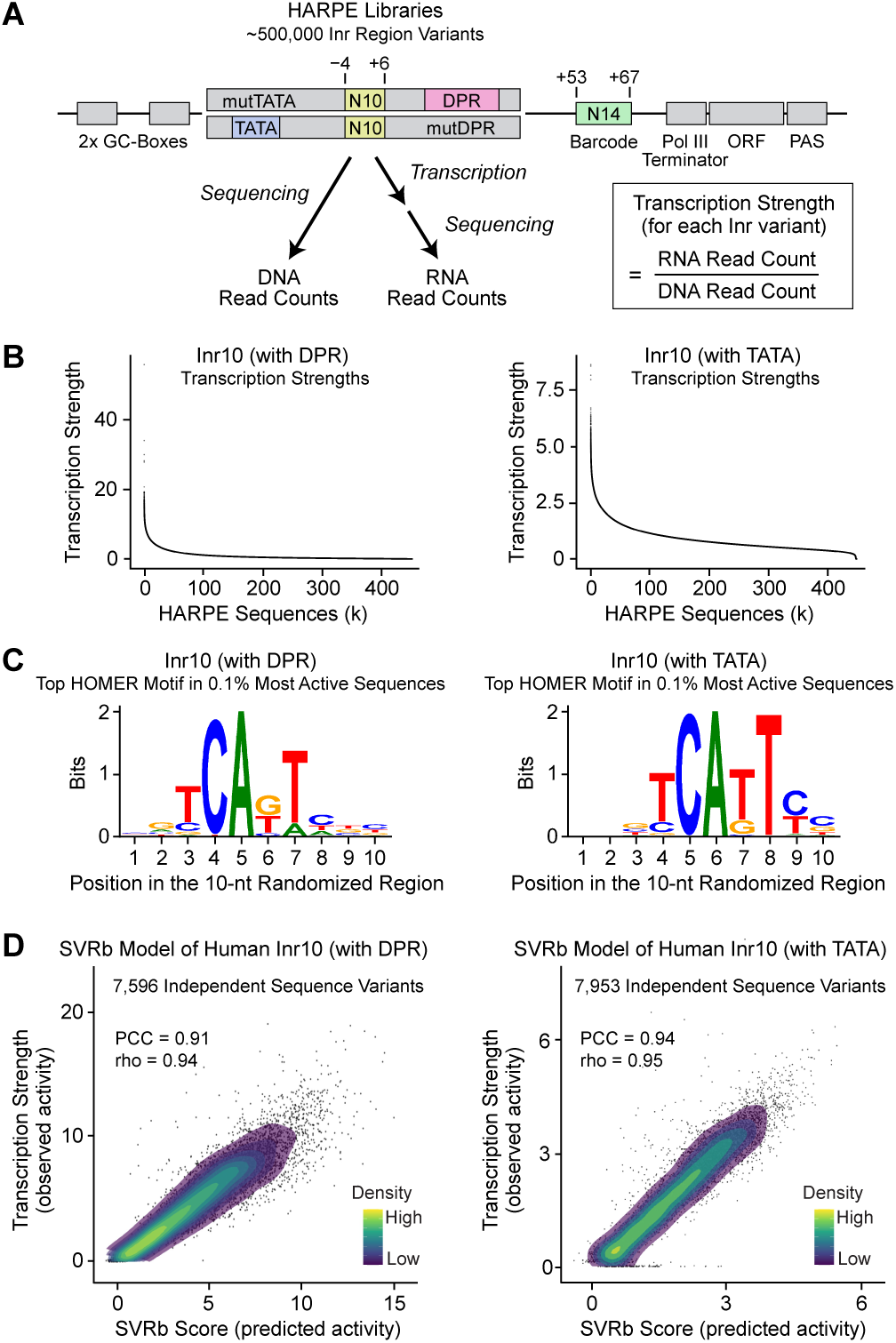
HARPE and SVR analyses of the human Inr in DPR- and TATA-driven promoter backbones. (*A*) Use of the HARPE method for the analysis of the human Inr region. The SCP1-based promoter backbones are identical except for the presence of a DPR (mutTATA) versus a TATA box (mutDPR). The 10-nt region from −4 to +6 (relative to the SCP1 +1 TSS; designated as “Inr10”) was randomized to generate approximately 500,000 Inr sequence variants in each promoter backbone. Each Inr sequence variant was identified with a corresponding 14-nt barcode from +53 to +67. (*B*) Most Inr sequence variants exhibit low transcriptional activity, but a small subset of the variants is highly active. Shown is the transcription strength (with biochemical assays, in order of decreasing activity) of each of the approximately 500,000 Inr variants with a DPR (left) or TATA box (right). (*C*) HOMER analysis of the 0.1% most active Inr variants (in biochemical assays) with a DPR (left) or TATA box (right) reveals a distinct motif that closely resembles the *Drosophila* Inr. Shown is the web logo of the top HOMER motif. (*D*) The SVRb (trained with biochemical data) models of the human Inr accurately predict the transcription strengths of Inr sequence variants with a DPR (left) or TATA box (right). Each SVRb model was trained with 200,000 variants and then tested with approximately 7,000-8,000 independent variants (*i.e.*, not used for training) from the HARPE dataset. For each of the independent test sequences, the predicted SVRb score was compared with the observed transcription strength. (PCC) Pearson’s correlation coefficient, (rho) Spearman’s rank correlation coefficient. SVR scores are relative (*i.e.*, not comparable between models) and are not equivalent to the observed transcription strengths. All panels show average transcription strengths from two independent biological replicates. The SVRb models were trained on the averaged values.

To provide a generally useful means of evaluating the human Inr, we used the HARPE data to train and optimize SVR models of the human Inr region (Supplemental Fig. S3), and obtained two SVR models, which we termed Inr10 (with DPR) SVRb and Inr10 (with TATA) SVRb. [Inr10 specifies the 10-nt randomized region; (with DPR) and (with TATA) denote the use of DPR- and TATA-driven core promoters, respectively; and SVRb indicates biochemical transcription data.] With these SVR models, we observed an excellent correlation between the observed transcription strengths of independent (*i.e.*, not used for training) test sequences and their predicted Inr activities (Fig. 1D). Thus, the Inr10 SVRb models of the human Inr provide accurate predictions of transcriptional activity.

We next investigated whether different libraries with a longer stretch of randomized Inr sequences would yield HARPE data and SVR models that are similar to those obtained with the Inr10 experiments. We therefore constructed and analyzed 14-nt randomized Inr region libraries (from −5 to +9 relative to the SCP +1 TSS) and found that the properties of the top 0.1% of variants and the resulting SVR models, termed Inr14 (with DPR) SVRb and Inr14 (with TATA) SVRb (Supplemental Fig. S4), are nearly identical to those obtained with the 10-nt Inr libraries (Supplemental Fig. S5). We also tested a third library in which both the DPR and the TATA box were mutated in the promoter backbone. This library exhibited much lower overall transcription levels but nevertheless yielded the same top HOMER motif (Supplemental Fig. S5; Heinz et al. 2010) and a good SVR model (Supplemental Fig. S4). Therefore, both the Inr10 and Inr14 SVRb models provide reliable predictions for the activity of the human Inr with the DPR as well as the TATA box.

Third, we tested whether transcription of the Inr region HARPE libraries in cells would yield results that are similar to those obtained in biochemical transcription reactions. We thus performed HARPE analyses by transfection of the 14-nt randomized Inr libraries into HeLa cells. The 0.1% most active cell-transcribed variants exhibited, as did the biochemical experiments (Supplemental Figs. S2,S5), a top HOMER motif that strongly resembles the *Drosophila* Inr (Supplemental Fig. S6). We generated cell-based Inr14 SVRc models (where SVRc indicates a cell-based SVR model; Supplemental Fig. S4) and found that the predictions of the SVRc models strongly correlate with those of their corresponding SVRb models (Fig. 2A,B). Thus, the biochemical and cell-based experiments lead to nearly identical conclusions about the human Inr.

**Figure 2.**
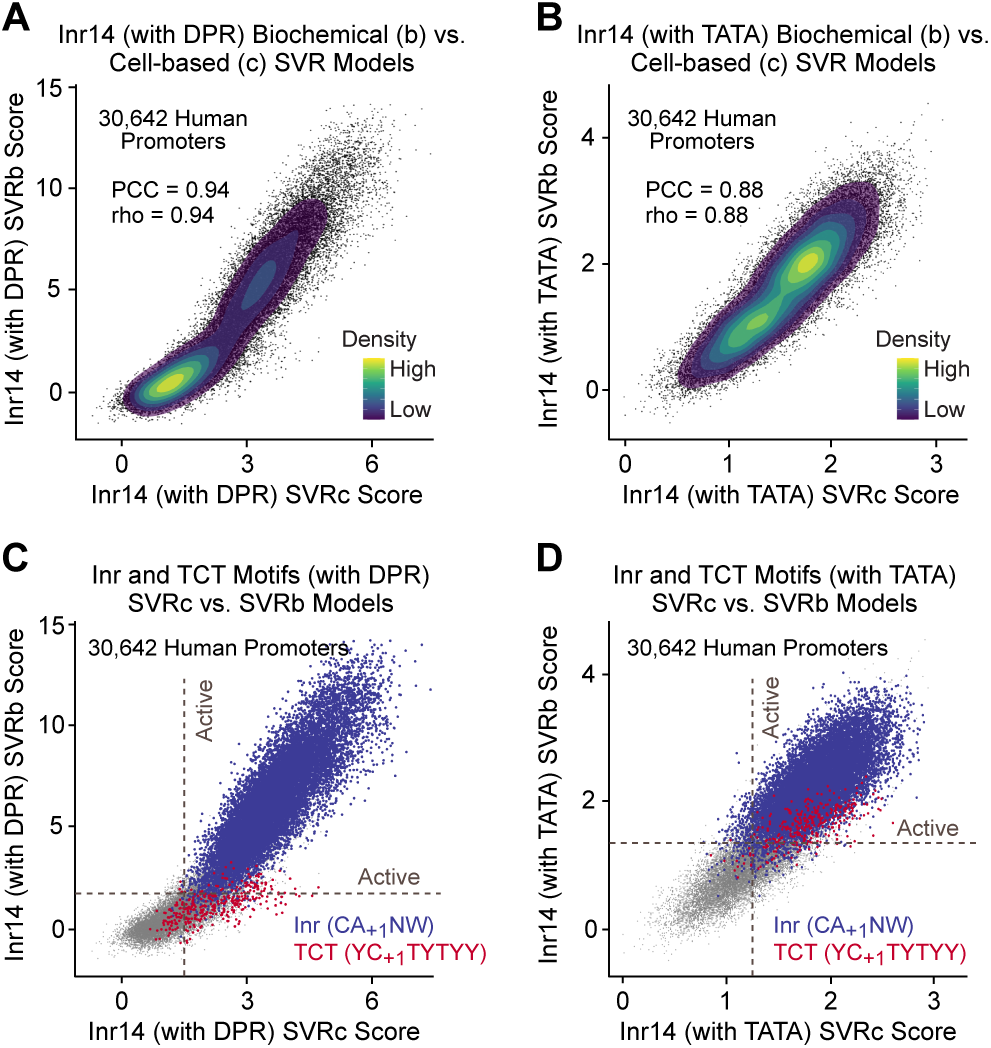
Analysis of the Inr and TCT motifs with biochemical (SVRb) and cell-based (SVRc) models of the human Inr. SVR models were generated with biochemical as well as cell-based HARPE data with a 14-nt randomized region (from −5 to +9 relative to the SCP +1 TSS; termed “Inr14”), which encompasses both the Inr and TCT motifs. (*A,B*) SVRc models of the human Inr with a DPR (*A*) or TATA box (*B*) show strong correlation with the corresponding SVRb models. SVRc and SVRb scores of 30,642 natural human promoters are compared. (PCC) Pearson’s correlation coefficient, (rho) Spearman’s rank correlation coefficient. (*C,D*) The TCT motif can be identified more distinctly with the cell-based Inr14 SVRc models than with the biochemical Inr14 SVRb models. SVR scores are shown for 30,642 natural human promoters, with sequences matching the Inr (CA_+1_NW) or TCT (YC_+1_TYTYY) motifs highlighted in blue or red, respectively. The red dots are plotted on top of the blue dots for visibility.

### Analysis of the Inr and TCT motif

To estimate the abundance of the Inr in natural human promoters, it was first necessary to determine the optimal SVR score thresholds for classifying sequences as active versus inactive. To this end, we carried out a performance assessment for each SVR model to identify the score thresholds at which the models are most accurate (Supplemental Figs. S7−S10). For example, with the Inr10 (with DPR) SVRb model, we found that 10-nt sequences with an SVR score ≥1.3 are predicted to be active as an Inr (Supplemental Fig. S7B). With the SVR score thresholds, we found that approximately 60% of natural human focused promoters contain an active Inr (Supplemental Fig. S11, Supplemental Table 2). [In these analyses, we examined focused promoters in which transcription initiates at a single site or in a narrow cluster of sites.] This observation is comparable to a recent estimate of ∼40-56% based on a consensus sequence analysis (Vo ngoc et al. 2017b). In addition, the different SVR models of the Inr (with a DPR, TATA box, or neither element) mostly agree on their predictions of which sequences are active Inr elements (Supplemental Fig. S11). These findings provide an independent revised assessment of the high abundance of the Inr in human promoters.

We additionally examined the shared features of the Inr sequences in natural promoters that are predicted to be active. To this end, we used the Inr10 SVRb models to generate sequence logos of all of the predicted active Inr sequences in human GM12878 cells, and identified a minimal CA_+1_NW motif (Supplemental Fig. S11E,F), which resembles the BBCA_+1_BW consensus that was based on the analysis of natural promoters and estimated to be present in about 40-56% of human promoters (Vo ngoc et al. 2017b). The CA_+1_NW consensus serves as a minimal Inr reference sequence, but, unlike the SVR models, CA_+1_NW does not provide any quantitative information on the transcriptional activity of any Inr that matches this motif. It is useful to note, however, that the Inr14 SVRb and SVRc models predict that most of the human Inr sequences with a match to the minimal CA_+1_NW Inr motif are active (Fig. 2C,D, Supplemental Fig. S12).

Importantly, in addition to the Inr, the randomized Inr region encompasses the overlapping but functionally distinct TCT motif (Supplemental Fig. S1), which is a rare but biologically important core promoter element that mediates the transcription of ribosomal protein genes and other genes involved in translation in animals (Parry et al. 2010; Wang et al. 2014). We therefore investigated whether the SVR models of the Inr region also recognize the activity of the TCT motif (YC_+1_TYTYY in humans; Parry et al. 2010). Unlike the Inr, the TCT motif is detected much more distinctly with the Inr14 SVRc models than with the Inr14 SVRb models (Fig. 2C,D, Supplemental Fig. S12). To determine the basis of this effect, we examined the observed transcriptional activities of TCT sequences in the HARPE assays and found that TCT sequences are much more active in cells than in biochemical experiments (Supplemental Fig. S13). Hence, the biochemical transcription extracts weakly detect the activity of TCT-dependent promoters. In contrast, the cell-based HARPE experiments effectively incorporate the activities of both the Inr and TCT motif. Notably, most of the identified TCT motifs are predicted to be active with the SVRc models (Fig. 2C,D, Supplemental Fig. S12). In addition, consistent with previous analyses of the TCT motif (Parry et al. 2010; Wang et al. 2014), a large fraction of the TCT-containing promoters are associated with ribosomal protein genes or genes that encode proteins that are involved in translation (Supplemental Table 3A,B). Thus, the Inr14 SVRc models can be used to predict the activity of both the Inr and the TCT motif in humans. It should also be noted that the Inr element alone, separately from the TCT motif, could be analyzed with the SVRb models, which provide strong predictions of Inr activity but only weakly detect the TCT motif.

*An Inr variant that functions specifically with the TATA box* We next compared the properties of the Inr in a DPR-driven versus a TATA-driven promoter. By using the Inr10 (with DPR) and Inr10 (with TATA) SVRb models, we determined the predicted Inr activities of all of the 1,048,576 possible 10-nt sequences (Supplemental Fig. S14A). This analysis revealed three Inr variants that differ in the positioning of the core Inr CAKT motif within the 10-nt randomized region. We then examined the occurrence of the three Inr variants in natural promoters and found that they are conserved from *Drosophila* to humans, as seen in human GM12878 cells (Fig. 3A), human K562 cells (Supplemental Fig. S14B), and *Drosophila* embryos (Fig. 3B). In the natural promoters, the A nucleotide in the core CAKT Inr motif is located at different positions (A+1, A+2, or A+3) relative to the observed +1 TSS. In these analyses, we used datasets that were generated by different methods to ensure that the TSS positions are not biased to any peculiarities of a single method. Hence, the TSSs in the GM12878 cells were identified by the GRO-cap method (Kruesi et al. 2013; Core et al. 2014), and the TSSs in the K562 cells and *Drosophila* embryos were determined with the csRNA-seq method (Duttke et al. 2019).

**Figure 3.**
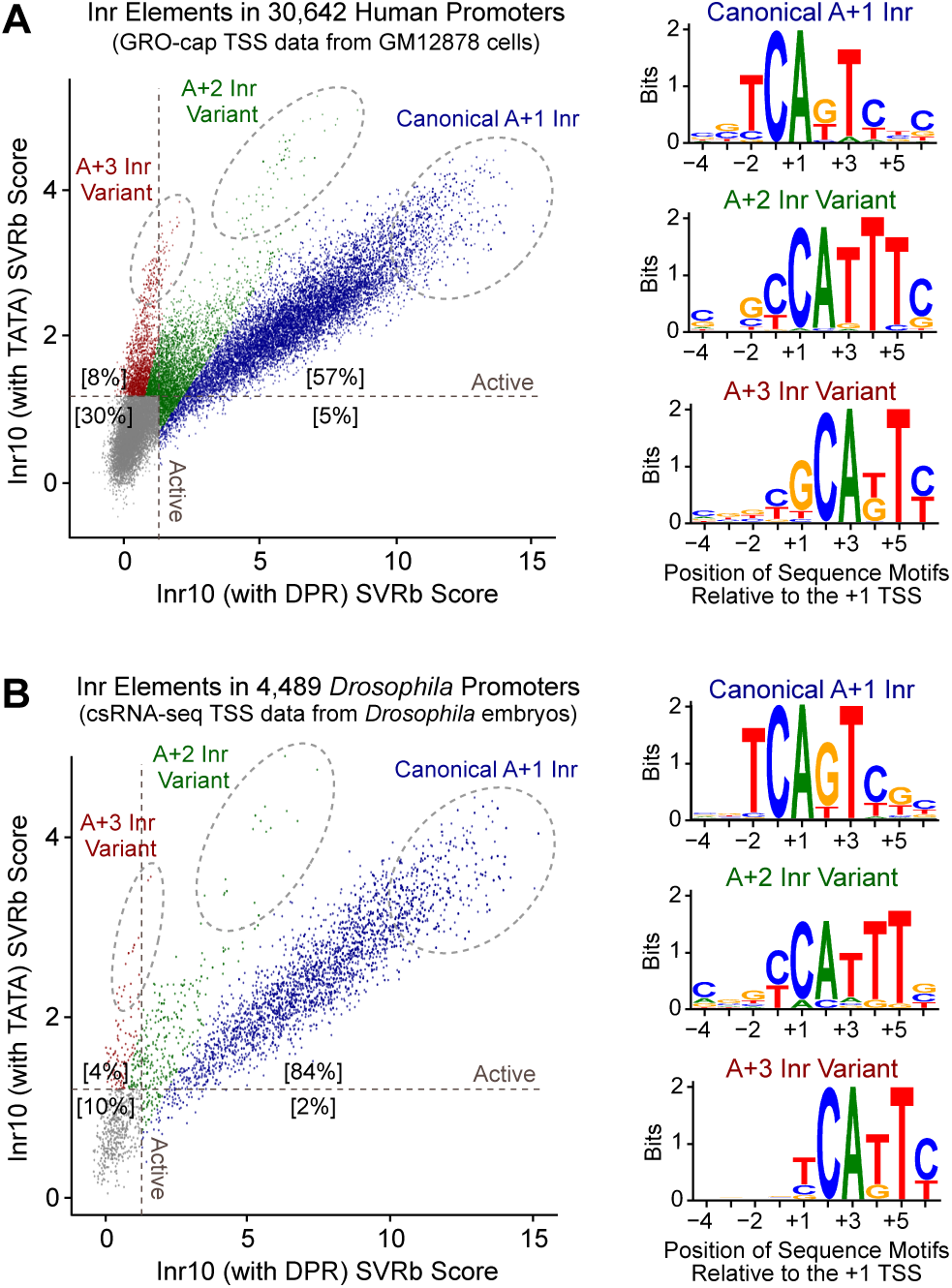
The positioning of the Inr sequence motif relative to the +1 TSS is different in DPR-driven promoters than in TATA-driven promoters. (*A*) Comparison of the predicted Inr activities of 30,642 natural human promoters with the Inr (with DPR) SVR model versus the Inr (with TATA) SVR model reveals three distinct variants, which are labeled according to the position of the A nt in the core CAKT Inr motif relative to the observed +1 TSS. The web logos are based on the sequences in the highlighted dashed ovals. (*B*) The three Inr variants are conserved from *Drosophila* to humans. Comparison of the predicted Inr activities of 4,489 natural *Drosophila* promoters with the Inr (with DPR) SVR model versus Inr (with TATA) SVR model reveals the same three Inr variants that are seen in human promoters.

The predicted activities of these three Inr variants differ in a DPR- versus a TATA-driven promoter (Fig. 3, Supplemental Fig. S14). With the Inr10 (with DPR) SVRb model, the canonical A+1 Inr is most active, the A+2 Inr is moderately active, and the A+3 Inr is inactive or very weakly active. This preference is consistent with the previously observed spacing requirement between the Inr and DPR (Burke and Kadonaga 1997; Kutach and Kadonaga 2000; Vo ngoc et al. 2020; Vo ngoc et al. 2023). In contrast, with the Inr10 (with TATA) SVRb model, the A+1, A+2, and A+3 Inr variants are predicted to be of comparable activity. The A+3 variant exhibits distinctly different properties with the TATA box relative to the DPR.

To examine the Inr variants from a different perspective, we compared their occurrence in natural TATA- versus DPR-containing promoters (Supplemental Fig. S15). Strikingly, we found that an active A+3 Inr is present in 8.3% (368/4,448) of TATA-containing human promoters, but only in 0.1% (10/7,630) of DPR-containing human promoters. Even more notably, an active A+3 Inr is present in 4.6% (46/1,009) of TATA-containing *Drosophila* promoters and is completely absent (0/3,070) in the DPR-containing *Drosophila* promoters. These findings collectively indicate that the A+3 Inr variant functions specifically with the TATA box relative to the DPR.

### A strong TATA box can drive transcription initiation in the absence of an Inr or DPR

By using the SVR models of the human Inr in conjunction with our previously generated models of the human TATA box (Vo ngoc et al. 2020), human DPR (Vo ngoc et al. 2020), and *Drosophila* DPR (Vo ngoc et al. 2023), we examined the cooccurrence of these elements in natural human and *Drosophila* promoters. A key feature of these analyses is that they provide quantitative genome-wide assessments rather than anecdotal observations of the relationships between the elements. When the promoters were ranked according to their Inr SVR scores, we observed a striking inverse relationship between predicted Inr and TATA activities (Fig. 4A,B, Supplemental Fig. S16). The duality between the TATA and DPR (see, for example: Willy et al. 2000; Hsu et al. 2008; Vo ngoc et al. 2020) can also be clearly seen in the heat maps that are ranked by the DPR SVR scores or the TATA SVR scores (Supplemental Fig. S16).

**Figure 4.**
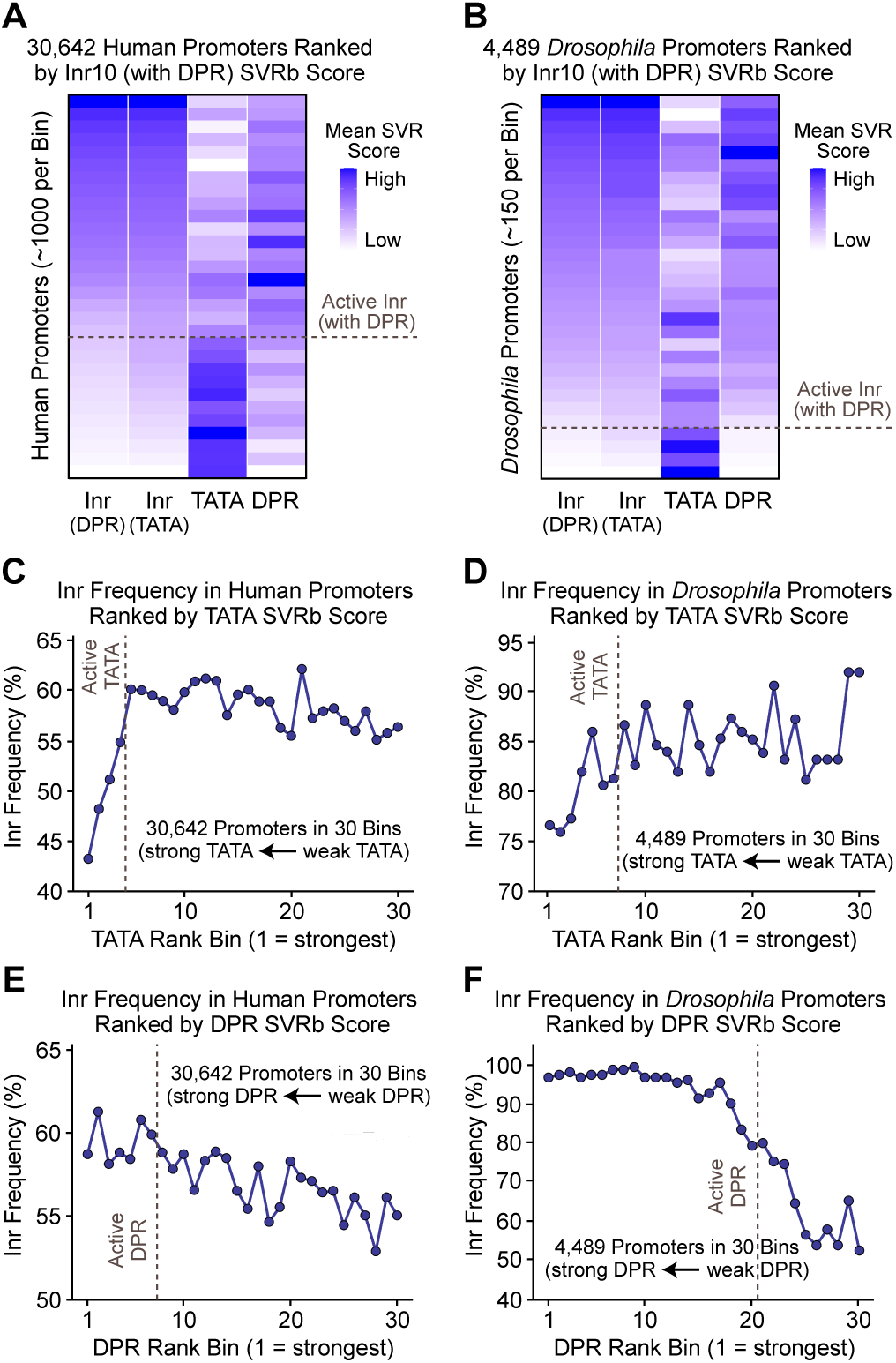
Predicted Inr activity positively correlates with predicted DPR activity and negatively correlates with predicted TATA box activity. Natural promoters were ranked according to their human Inr (with DPR) scores and divided into 30 bins of approximately equivalent size. For each bin, the mean SVR scores for human Inr (with DPR), human Inr (with TATA), human TATA box (Vo ngoc et al. 2020), and human or *Drosophila* DPR (Vo ngoc et al. 2020, 2023) were determined. (*A,B*) Positive correlation between the DPR and Inr and negative correlation between the TATA box and Inr in both humans (*A*) and *Drosophila* (*B*). (*C,D*) The predicted frequency of active Inr elements sharply declines with the predicted strength of active TATA motifs in human (*C*) and *Drosophila* (*D*) promoters. (*E,F*) The predicted frequency of active Inr elements increases with the predicted strength of DPR motifs in human (*E*) and *Drosophila* (*F*) promoters. In *C–F*, active Inr elements were defined to be those that are predicted to be active with both the Inr10 (with DPR) and Inr10 (with TATA) SVR models.

Further analysis of the Inr and TATA revealed that the predicted frequency of active Inr elements sharply declines with the predicted strength of active TATA motifs (Fig. 4C,D, Supplemental Fig. S17A,B). This effect was seen in two different human cell lines as well as in *Drosophila* embryos and S2 cells. In contrast, the predicted Inr and DPR activities are positively correlated (Fig. 4A,B, Supplemental Fig. S16), with the predicted frequency of active Inr elements increasing with the predicted strength of DPR motifs (Fig. 4E,F, Supplemental Fig. S17C,D). The positive correlation between the Inr and DPR is consistent with the synergy between these elements (Burke and Kadonaga 1996). A negative correlation between the TATA and Inr was previously observed, as assessed with an Inr consensus sequence (Vo ngoc et al. 2020). The SVR models of the Inr further revealed the sharp decline of the Inr frequency with increasing strength of active TATA elements.

To examine the interrelationships between the TATA, Inr, and DPR elements, we carried out permutation testing with natural human and *Drosophila* promoters (Supplemental Fig. S18). This analysis revealed two significantly enriched core promoter types – those containing only a TATA box [TATA+, Inr−, DPR−] and those containing both an Inr and DPR [TATA−, Inr+, DPR+]. The enrichment of the [TATA−, Inr+, DPR+] promoters is in accord with the previously noted synergy between the Inr and DPR as well as the cooccurrence of these elements (Kutach and Kadonaga 2000; Vo ngoc et al. 2020). The enrichment of the [TATA+, Inr−, DPR−] promoters is consistent with the inverse correlation between the TATA and Inr (Fig. 4, Supplemental Figs. S16,S17) and the duality between the TATA box and DPR. These findings collectively indicate that the Inr appears to play a more important role in DPR-driven transcription than in TATA-driven transcription and that a strong TATA box can drive transcription initiation in the absence of an Inr or DPR.

It is also notable that we observed a much larger dynamic range of Inr activity with a DPR than with a TATA box (Supplemental Fig. S19). With a DPR-driven promoter, the strongest Inr variants are ∼100-fold more active than the median inactive sequence, whereas with a TATA-driven promoter, the strongest Inr variants are only ∼10-fold more active than the median inactive sequence. These findings further indicate that there is a stronger functional relationship between the Inr and DPR than between the Inr and TATA. Thus, the high throughput HARPE data, the SVR models of the Inr, TATA, and DPR, and the genome-wide human and *Drosophila* promoter analyses collectively lead to a model in which there is a strict and synergistic interaction between the Inr and DPR, a duality between the TATA and DPR, and a flexible and sometimes independent function of the TATA box in relation to the Inr.

### Analysis and perspectives

The Inr is a widely-used transcriptional element that is present in about 60% of human promoters. Here, we combined high-throughput transcriptional analyses with machine learning to generate highly predictive models of the human Inr. The SVR models are effective at predicting active Inr elements, and it is thus reasonable to ask, anthropomorphically, “What are the SVR models thinking?” To address this question, we first determined the predicted Inr activity of all of the 1,048,576 possible 10-nt sequences and identified the predicted optimal Inr motif (Supplemental Fig. S20A). Second, to evaluate the importance of each position within the Inr region, we randomized the nucleotides at each position and measured the resulting decrease in model performance (Supplemental Fig. S20B). This analysis revealed that the positions corresponding to the minimal CANW Inr sequence are the most important for Inr activity, but also that all four nucleotides in this motif contribute to Inr function. Third, we evaluated the effect of every nucleotide substitution at each position across all 10-nt sequences, and identified position- and nucleotide-specific effects on predicted transcriptional activity (Supplemental Fig. S20C). Collectively, these model interpretation analyses uncover the key sequence features that determine Inr activity.

There are different types of core promoters that have different properties based on the presence or absence of motifs such as the TATA box and DPR. For instance, in *Drosophila*, TATA-specific and DPR-specific transcriptional enhancers have been identified (Butler and Kadonaga 2001; Juven-Gershon et al. 2008). The SVR models have enabled us to gain a better understanding of the relationships between the TATA box, Inr, and DPR elements, and have thus provided new insights into the nature of the different types of core promoters. In this regard, we further investigated the TCT motif. Because the TCT motif is rare and overlaps the Inr, we were not able to generate a TCT-specific SVR model. Therefore, for the analysis of the TCT motif, we used sequences that match the human TCT consensus (YC_+1_TYTYY), and found that the relation between the TATA box and TCT motif in humans is, strikingly, opposite from that seen in *Drosophila* (Supplemental Fig. S21). Specifically, we observed that the TATA box is overrepresented in human TCT promoters and underrepresented in *Drosophila* TCT promoters. We likewise found that the TCT motif is enriched in human TATA-containing promoters and depleted in *Drosophila* TATA-containing promoters. These findings indicate that the properties of the TCT motif are different in humans than in *Drosophila*. This point should be taken into consideration in the analysis of the TCT motif in humans.

It is also important to consider whether different types of core promoter elements are associated with specific biological functions. For instance, the TCT motif is used by the ribosomal protein gene network (Parry et al. 2010). To explore the possible biological functions that may be associated with the two enriched core promoter types (TATA-only and Inr-DPR; Supplemental Fig. S18), we performed gene ontology (GO) analyses with human promoters (Supplemental Table 4A,B). We found that genes associated with TATA-only promoters are enriched for many functions related to signaling, immune response, and stress response, which suggests that genes that mediate responses to changing conditions may frequently employ TATA-driven transcription. In contrast, the Inr-DPR promoters did not yield any significantly enriched GO terms. These results suggest that there are different biological functions of TATA-only and Inr-DPR core promoters in human gene regulation.

Somewhat remarkably, we found that the optimal human Inr, TCA_+1_KTY, is essentially identical to the *Drosophila* Inr consensus (see, for example: Hultmark et al. 1986; Ohler et al. 2002, FitzGerald et al. 2006). Similarly, in Vo ngoc et al. (2023), we found that the DPR consensus sequence is nearly the same in humans and *Drosophila*. It thus appears that the Inr and DPR motifs have remained largely unchanged over the ∼700 million years of species divergence time between *Drosophila* and humans (Kumar et al. 2022), despite differences in promoter G+C content (Deaton and Bird 2011).

Thus, in conclusion, we were able to generate SVR models of the human Inr and to gain key insights into its occurrence as well as its relation to other core promoter elements at the genome-wide level. There remains, however, much to be learned. In the future, we will seek to identify the DNA motifs and protein factors that drive transcription from the roughly 30% of human promoters that lack any of the known core promoter elements. We also plan to integrate our understanding of the core promoter with the functions of promoter- and enhancer-binding proteins. In this manner, it may be possible to uncover new unified perspectives into the specific molecular interactions that mediate the regulation of transcription by RNA polymerase II.

## Materials and Methods

### HARPE analysis of the Inr

The HARPE plasmid libraries for the Inr were generated as described in Vo ngoc et al. (2020). The HARPE vector and SCP1-based promoter backbones, which include constructs with mutations in the DPR and/or the TATA box, are identical to those used previously (Vo ngoc et al. 2020, 2023). For HARPE analysis of the Inr, the 14-nt region from −5 to +9 and the 10-nt region from −4 to +6 (relative to the SCP1 +1 TSS) were randomized to generate ∼500,000 Inr sequence variants within SCP1 promoter backbones containing a DPR (mutTATA), a TATA box (mutDPR), or neither element (mutTATA, mutDPR). In vitro transcription with HeLa nuclear extracts and transfection into HeLa cells were performed as described in Vo ngoc et al. (2020). Sample and data processing were performed as described in Vo ngoc et al. (2020, 2023). Illumina sequencing of NGS PCR amplicons was performed on NovaSeq 6000 or NovaSeq X Plus sequencing systems at the IGM Genomics Center, University of California, San Diego. Additional information on HARPE library generation, transcription, RNA processing, sequencing, and data analysis is provided in the Supplemental Material.

### Generation and use of the SVR models

Machine learning analyses were performed as described in Vo ngoc et al. (2020, 2023), except that we used functions from the Python package scikit-learn (version 1.3.0; Pedregosa et al. 2011) instead of the R package e1071 (version 1.7-2). Each SVR model was trained with 200,000 HARPE variants and then evaluated with ∼7,000-8,000 independent variants (i.e., not used for training). Grid search was used to optimize the hyperparameters cost and gamma, and cross validation was carried out with two independent test sets. Additional information on model training and optimization is provided in the Supplemental Material. We used SVR models of the human Inr (this study), human TATA box (“SVRTATA”; Vo ngoc et al. 2020; GSE139635), human DPR (“SVRb”; Vo ngoc et al. 2020; GSE139635, which is the same as “SVRh”; Vo ngoc et al. 2023; GSE225570) and *Drosophila* DPR (“SVRd”, Vo ngoc et al. 2023; GSE225570).

## Competing Interest Statement

The authors declare no competing interests.

## Acknowledgments

We thank Grisel Cruz-Becerra and George Kassavetis for critical reading of the manuscript. J.T.K. is the Amylin Chair in the Life Sciences. This publication includes data generated at the UC San Diego IGM Genomics Center utilizing Illumina NovaSeq 6000 and NovaSeq X Plus systems that were purchased with funding from National Institutes of Health SIG grant (#S10 OD026929). This work used Expanse CPU at the San Diego Supercomputer Center through the Advanced Cyberinfrastructure Coordination Ecosystem: Services & Support (ACCESS) program (project BIO230152), which is supported by NSF grants 2138259, 2138286, 2138307, 2137603, and 2138296. T.E.R.-C. received support from NIH grant T32 GM133351 to UCSD Biological Sciences Pathways in Biological Sciences (PiBS) Training Program and NSF Graduate Research Fellowship (GRFP) DGE-2545911. This work was supported by NIH grant R35 GM118060 to J.T.K. This paper was typeset with the bioRxiv word template by @Chrelli: www.github.com/chrelli/bio-Rxiv-word-template.

## Author Contributions

T.E.R.-C., L.V.n., and J.T.K. initially conceived the project. T.E.R.-C. performed most of the laboratory experiments as well as all computational analyses. L.V.n, C.M., and K.E.G. performed some experiments. T.E.R.-C. and J.T.K. prepared the figures and wrote the manuscript. J.T.K. oversaw the overall execution of this work.

## SUPPLEMENTAL MATERIAL

**Supplemental Figure S1.**
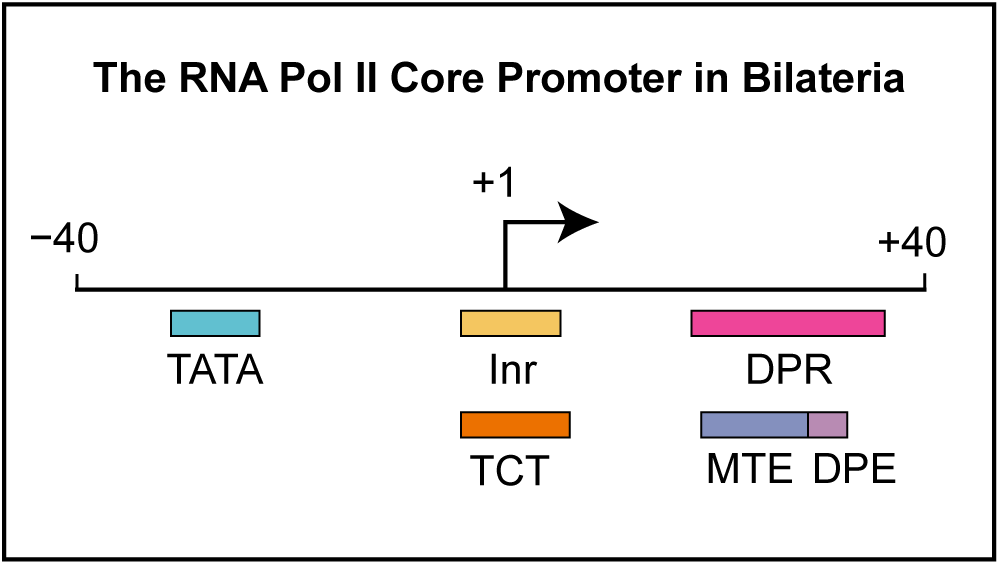
The RNA polymerase II core promoter in bilateria. This diagram depicts the DNA sequence motifs that are discussed in the text. The DPR is a unified combination of the MTE and DPE motifs. Because the DPR includes the DPE, we use the term “DPR” when referring to previous work on the DPE or DPR. The Inr and TCT motifs are overlapping but functionally distinct. The drawing is approximately to scale.

**Supplemental Figure S2.**
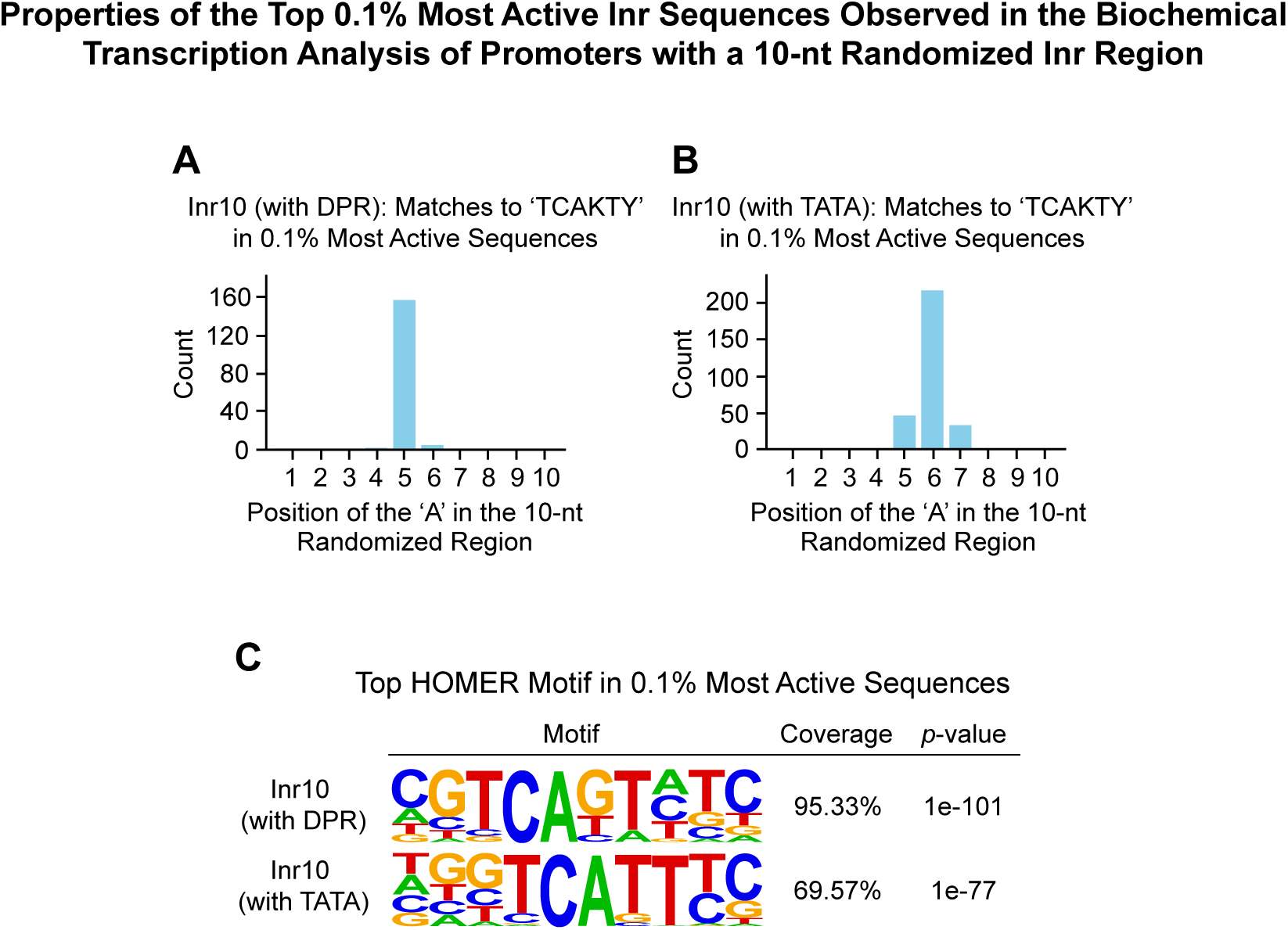
Properties of the top 0.1% most active Inr sequences observed in the biochemical transcription analysis of promoters with a 10-nt randomized Inr region. (*A*) In DPR-driven transcription (with a mutant TATA box), the ‘A’ in the TCAKTY Inr motif is sharply positioned at the 5th nt in the 10-nt randomized region. This position corresponds to the A+1 TSS in SCP1, in which the DPR is located from +17 to +35. This figure depicts the distribution of the ‘A’ nts in sequences with a perfect match to TCAKTY (*n* = 162) in the top 0.1% most active Inr (with DPR) sequences (*n* = 450). (*B*) In TATA-driven transcription (with a mutant DPR), the preferred position of the ‘A’ in the TCAKTY Inr motif is the 6th nt in the 10-nt randomized region. For reference, the upstream ‘T’ nucleotide in the SCP1 TATA box is located at the −32 position relative to the peak +1 TSS at the 6th nt. This figure shows the distribution of the ‘A’ nts in sequences with a perfect match to TCAKTY (*n* = 293) in the top 0.1% most active Inr (with TATA) sequences (*n* = 447) (*C*) HOMER analysis of the top 0.1% most active Inr variants with a DPR (top) or TATA box (bottom) reveals a distinct motif that closely resembles the *Drosophila* Inr. Shown are the top HOMER motifs.

**Supplemental Figure S3.**
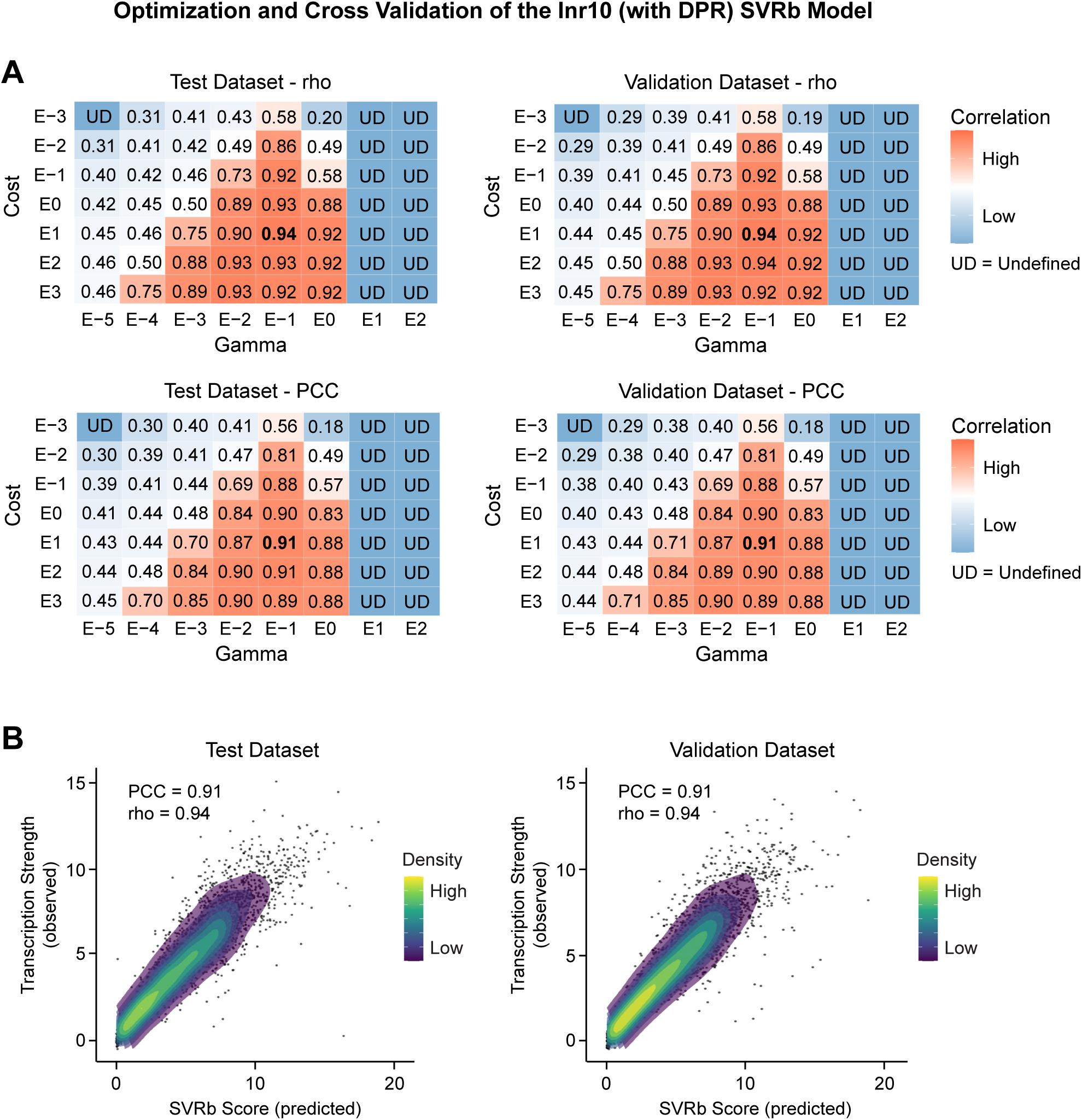
Optimization and cross validation of the Inr10 (with DPR) SVRb model. (*A*) Grid search results across different values of the cost and gamma hyperparameters, which control error tolerance and the influence of individual training sequences, respectively. Each matrix displays the values of Spearman’s rank correlation coefficient (rho; upper panels) or Pearson’s correlation coefficient (PCC; lower panels) for SVR models that were generated with the indicated values of cost and gamma with the test dataset (left panels) or the validation dataset (right panels), which are separate halves of the 7,596 independent test sequences shown in Fig. 1D (left). Undefined (UD) correlation is observed when the prediction of a model is constant regardless of the sequence. The hyperparameter values that were selected for the Inr10 (with DPR) SVR model are cost = 10 = E1 and gamma = 0.1 = E−1. (*B*) Comparison of observed transcription strengths (HARPE data) and predicted transcription strengths [Inr10 (with DPR) SVRb scores] with the test dataset (left) and validation dataset (right). All of the other SVR models were optimized and cross validated in the same manner.

**Supplemental Figure S4.**
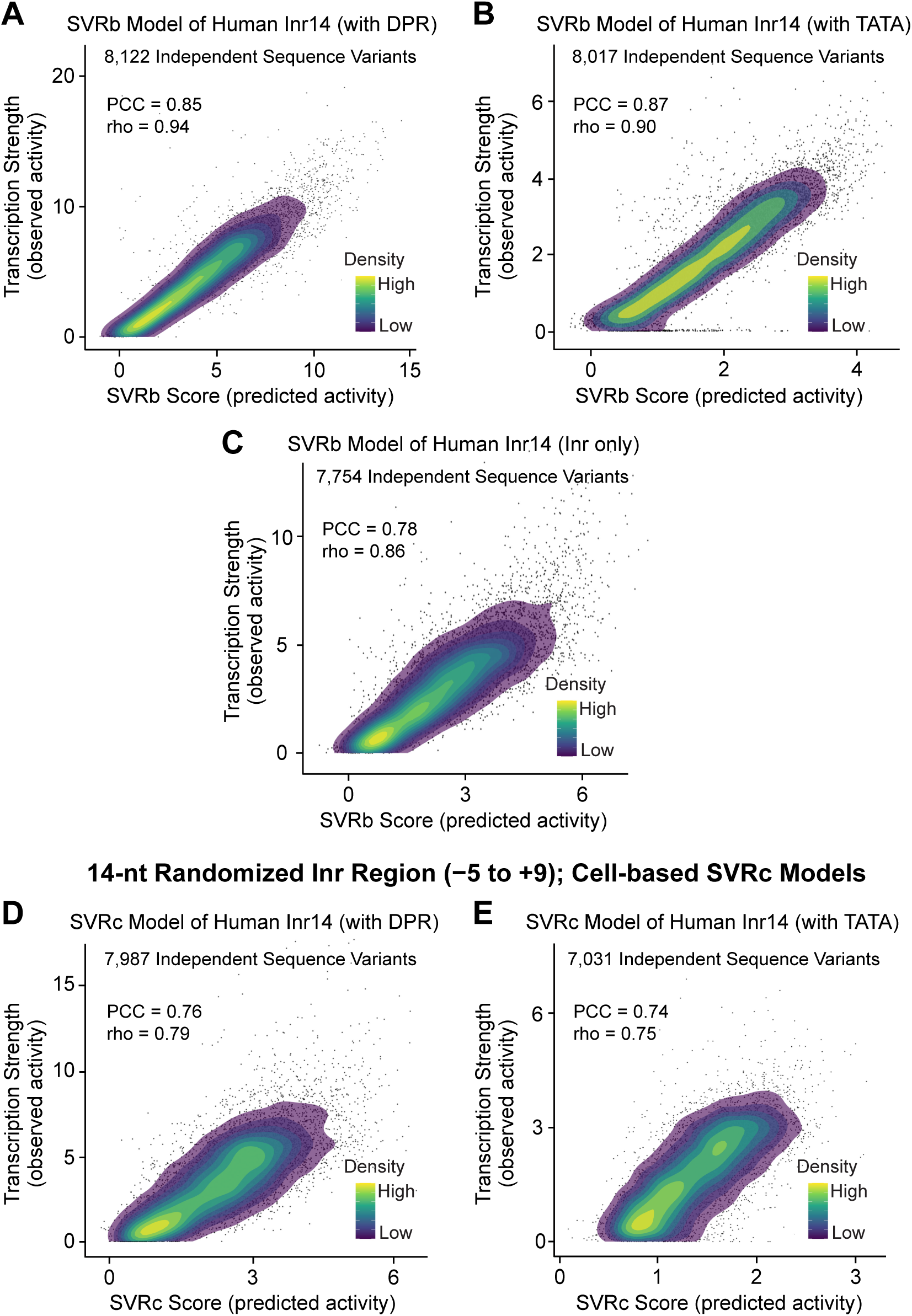
Biochemical (SVRb) and cell-based (SVRc) models of the human Inr accurately predict the transcription strengths of Inr sequence variants with a DPR, TATA box, or neither element. SVR models were trained on biochemical (*A,B,C*) or cell-based (*D,E*) HARPE data with a 14-nt randomized Inr region (−5 to +9 relative to the SCP +1 TSS; “Inr14”) in a DPR-driven (*A,D*), TATA-driven (*B,E*), or Inr-only (*C*) promoter backbone. Each SVR model was trained with 200,000 variants and then tested with approximately 7,000-8,000 independent variants (*i.e.*, not used for training) from the HARPE dataset. For each of the independent test sequences, the predicted SVR score was compared with the observed transcription strength. (PCC) Pearson’s correlation coefficient, (rho) Spearman’s rank correlation coefficient. SVR scores are relative (*i.e.*, not comparable between models) and are not equivalent to the observed transcription strengths. Shown are the average transcription strengths from two (*A,B,D,E*) or three (*C*) independent biological replicates. The SVR models were trained on the averaged values.

**Supplemental Figure S5.**
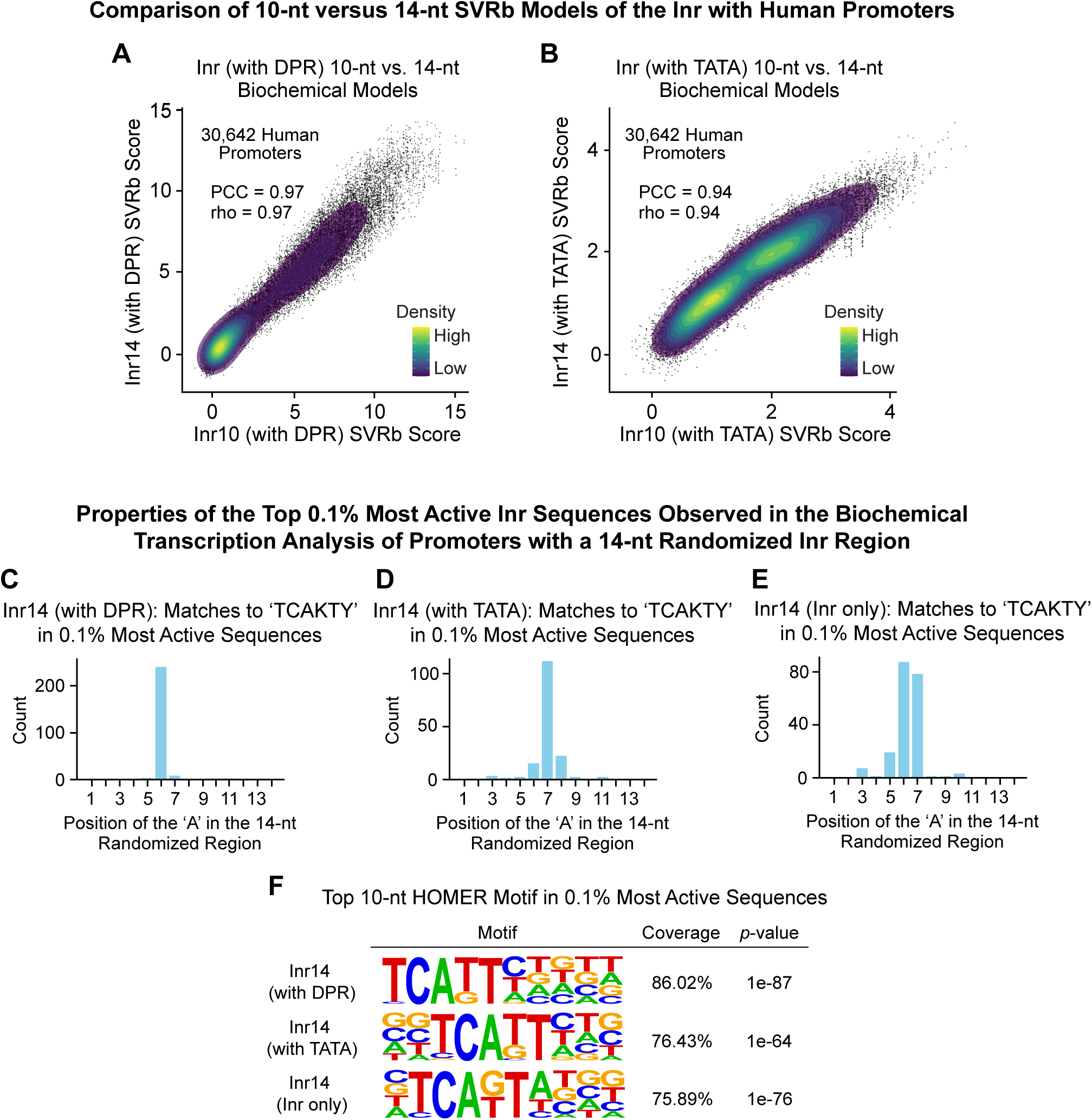
Properties of the human Inr based on the biochemical transcription analysis of promoters with a 14-nt randomized Inr region. (*A,B*) Inr14 SVRb models with a DPR-driven promoter (*A*) or a TATA box-driven promoter (*B*) show strong correlation with their corresponding Inr10 SVRb models. Inr10 and Inr14 SVRb scores of 30,642 natural human promoters are compared. (PCC) Pearson’s correlation coefficient, (rho) Spearman’s rank correlation coefficient. (*C,D,E,F*) Properties of the top 0.1% most active Inr sequences observed in biochemical transcription analyses with a 14-nt randomized Inr region. (*C*) In DPR-driven transcription (with a mutant TATA box), the ‘A’ in the TCAKTY Inr motif is sharply positioned at the 6th nt in the 14-nt randomized region. This position corresponds to the A+1 TSS in SCP1, in which the DPR is located from +17 to +35. This figure depicts the distribution of the ‘A’ nts in sequences with a perfect match to TCAKTY (*n* = 251) in the top 0.1% most active Inr (with DPR) sequences (*n* = 465). (*D*) In TATA-driven transcription (with a mutant DPR), the preferred position of the ‘A’ in the TCAKTY Inr motif is the 7th nt in the 14-nt randomized region. For reference, the upstream ‘T’ nucleotide in the SCP1 TATA box is located at the −32 position relative to the peak +1 TSS at the 7th nt. This figure shows the distribution of the ‘A’ nts in sequences with a perfect match to TCAKTY (*n* = 159) in the top 0.1% most active Inr (with TATA) sequences (*n* = 314) (*E*) In the absence of a DPR or a TATA box, the ‘A’ in the TCAKTY Inr motif is most often positioned at the 6th or 7th nt in the 14-nt randomized region. These positions correspond to the +1 or +2 locations relative to the SCP1 +1 TSS. This figure shows the distribution of the ‘A’ nts in sequences with a perfect match to TCAKTY (*n* = 197) in the top 0.1% most active Inr (with TATA) sequences (*n* = 336). (*F*) HOMER analysis of the top 0.1% most active Inr variants with a DPR (top), TATA box (middle), or neither (bottom) reveals a distinct motif that closely resembles the *Drosophila* Inr. Shown are the top 10-nt HOMER motifs.

**Supplemental Figure S6.**
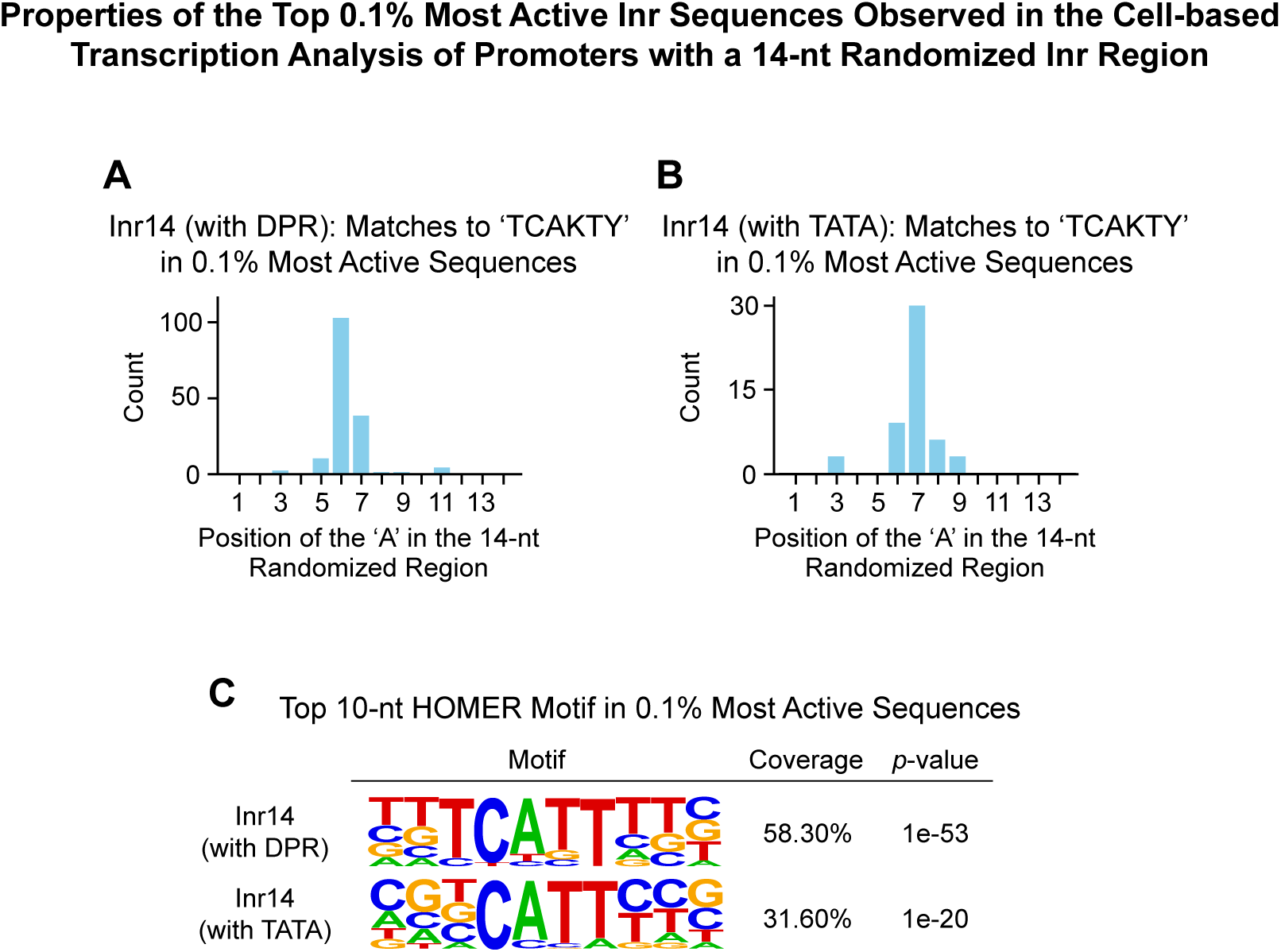
Properties of the top 0.1% most active Inr sequences observed in the cell-based transcription analysis of promoters with a 14-nt randomized Inr region. (*A*) In DPR-driven transcription (with a mutant TATA box), the preferred position of the ‘A’ in the TCAKTY Inr motif is 6th position in the 14-nt randomized region. This position corresponds to the A+1 TSS in SCP1, in which the DPR is located from +17 to +35. This figure depicts the distribution of the ‘A’ nts in sequences with a perfect match to TCAKTY (*n* = 158) in the top 0.1% most active Inr (with DPR) sequences (*n* = 446). (*B*) In TATA-driven transcription (with a mutant DPR), the preferred position of the ‘A’ in the TCAKTY Inr motif is the 7th nt in the 14-nt randomized region. For reference, the upstream ‘T’ nucleotide in the SCP1 TATA box is located at the −32 position relative to the peak +1 TSS at the 7th nt. This figure shows the distribution of the ‘A’ nts in sequences with a perfect match to TCAKTY (*n* = 51) in the top 0.1% most active Inr (with TATA) sequences (*n* = 307). (*C*) HOMER analysis of the top 0.1% most active Inr variants with a DPR (top) or TATA box (bottom) reveals a distinct motif that closely resembles the *Drosophila* Inr. Shown are the top 10-nt HOMER motifs.

**Supplemental Figure S7.**
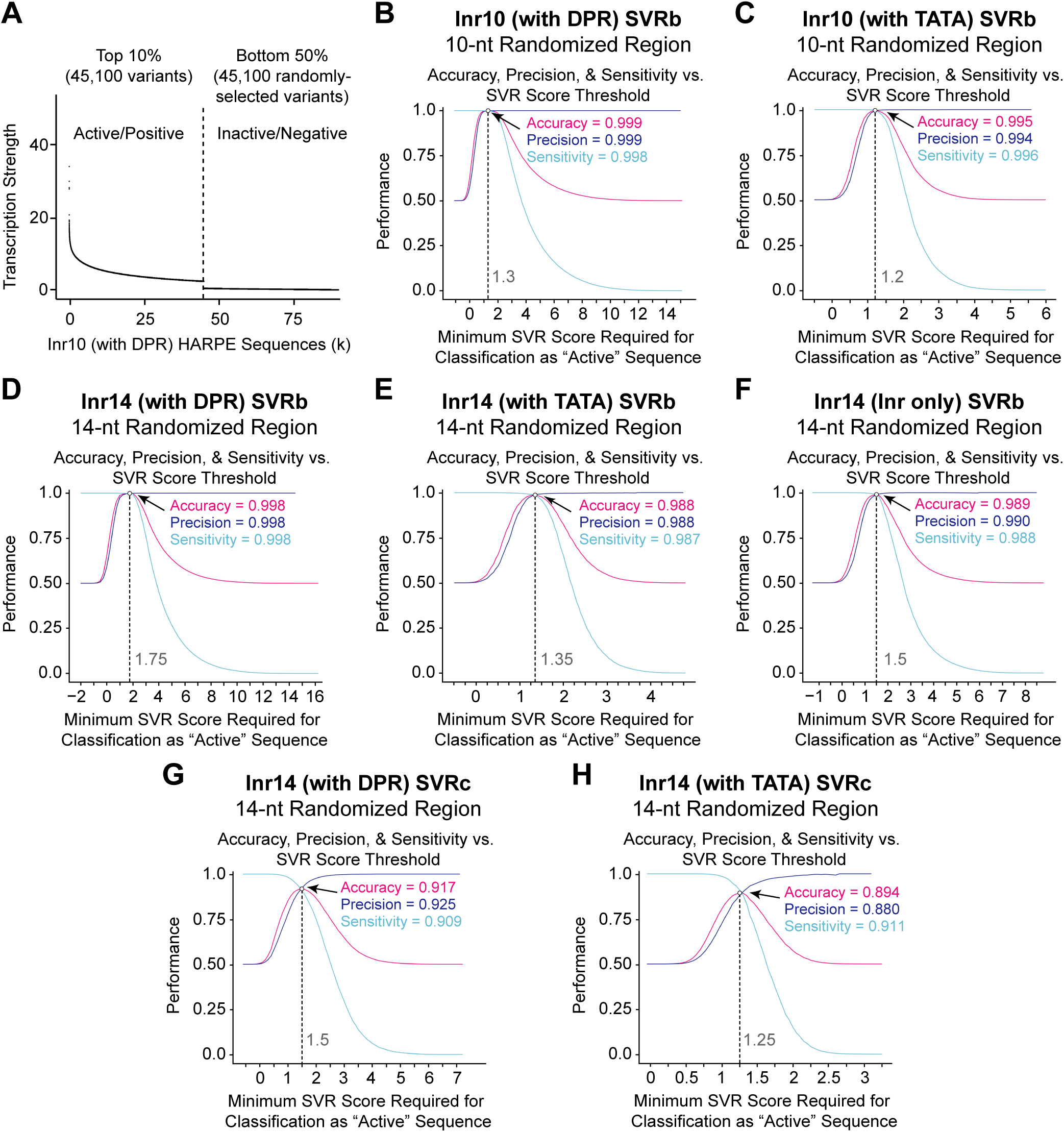
Determination of the optimal thresholds for distinguishing active versus inactive Inr elements via SVR performance assessment analyses of the accuracy, precision, and sensitivity versus the minimum SVR scores required for classification as an active sequence. (*A*) Selection of Inr10 (with DPR) HARPE sequence variants for the Inr10 (with DPR) SVRb performance assessment. The top 10% of variants were designated as active/positive for transcription, and an equal number of randomly-sampled sequences in the bottom 50% were designated as inactive/negative. Variants with intermediate activity were excluded. Shown are the transcription strengths (averaged across two independent HARPE replicates) of the selected variants. The same selection criteria were used for the other HARPE experiments. (*B–H*) Performance measures relative to the minimum SVR score required for a positive prediction. Performance was computed by counting true positives (TP), true negatives (TN), false positives (FP), and false negatives (FN). Accuracy [(TP+TN) / (TP+FP+TN+FN)] reflects how often SVR predictions are correct. Precision [TP / (TP + FP)] is the proportion of positive predictions that are correct. Sensitivity (or recall or true positive rate) [TP / (TP + FN)] is the proportion of transcriptionally active variants that are correctly predicted as positives. (*B*) Inr10 (with DPR) SVRb performance is optimal with 1.3 as the threshold for distinguishing active (SVR 2’ 1.3) versus inactive (SVR < 1.3) Inr sequences. (*C*) Inr10 (with TATA) SVRb performance is optimal with 1.2 as the threshold for distinguishing active (SVR 2’ 1.2) versus inactive (SVR < 1.2) Inr sequences. (*D*) Inr14 (with DPR) SVRb performance is optimal with 1.75 as the threshold for distinguishing active (SVR 2’ 1.75) versus inactive (SVR < 1.75) Inr sequences. (*E*) Inr14 (with TATA) SVRb performance is optimal with 1.35 as the threshold for distinguishing active (SVR 2’ 1.35) versus inactive (SVR < 1.35) Inr sequences. (*F*) Inr14 (Inr only) SVRb performance is optimal with 1.5 as the threshold for distinguishing active (SVR 2’ 1.5) versus inactive (SVR < 1.5) Inr sequences. (*G*) Inr14 (with DPR) SVRc performance is optimal with 1.5 as the threshold for distinguishing active (SVR 2’ 1.5) versus inactive (SVR < 1.5) Inr sequences. (*H*) Inr14 (with TATA) SVRc performance is optimal with 1.25 as the threshold for distinguishing active (SVR 2’ 1.25) versus inactive (SVR < 1.25) Inr sequences.

**Supplemental Figure S8.**
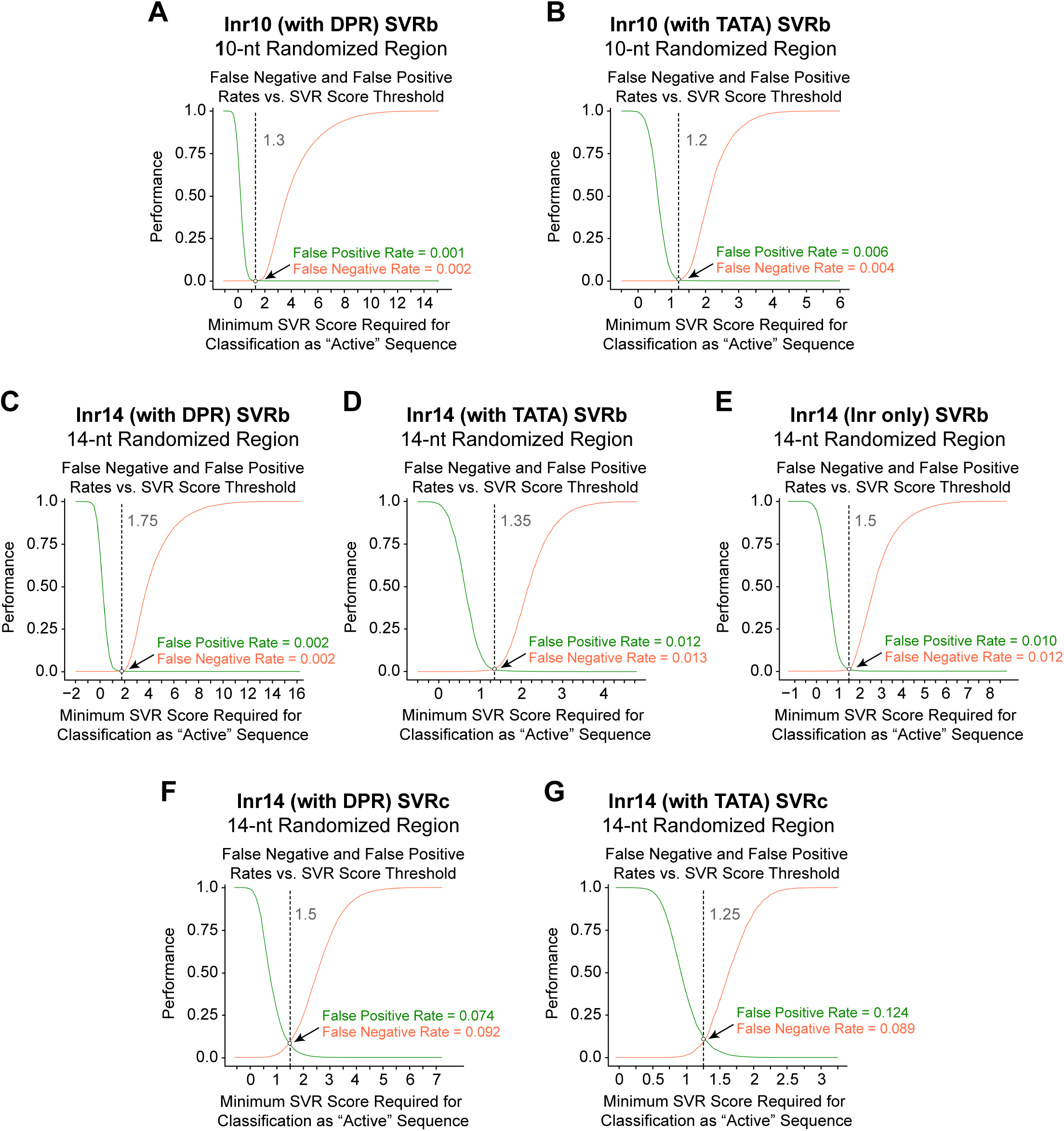
Determination of the optimal thresholds for distinguishing active versus inactive Inr elements via SVR performance assessment analyses of the false negative and false positive rates versus the minimum SVR scores required for classification as an active sequence. The HARPE sequence variants used for these performance assessments are the same as in Supplemental Fig. S7. (*A–G*) Performance measures relative to the minimum SVR score required for a positive prediction with the same SVR models described in Supplemental Fig. S7. Performance was computed by counting true positives (TP), true negatives (TN), false positives (FP), and false negatives (FN). False positive rate [FP / (FP + TN)] is the probability for an inactive sequence to be incorrectly predicted as positive. False negative rate [FN / (FN + TP)] = (1 − Sensitivity) is the probability for an active sequence to be incorrectly predicted as negative. The optimal SVR thresholds shown here are the same as those identified in Supplemental Fig. S7.

**Supplemental Figure S9.**
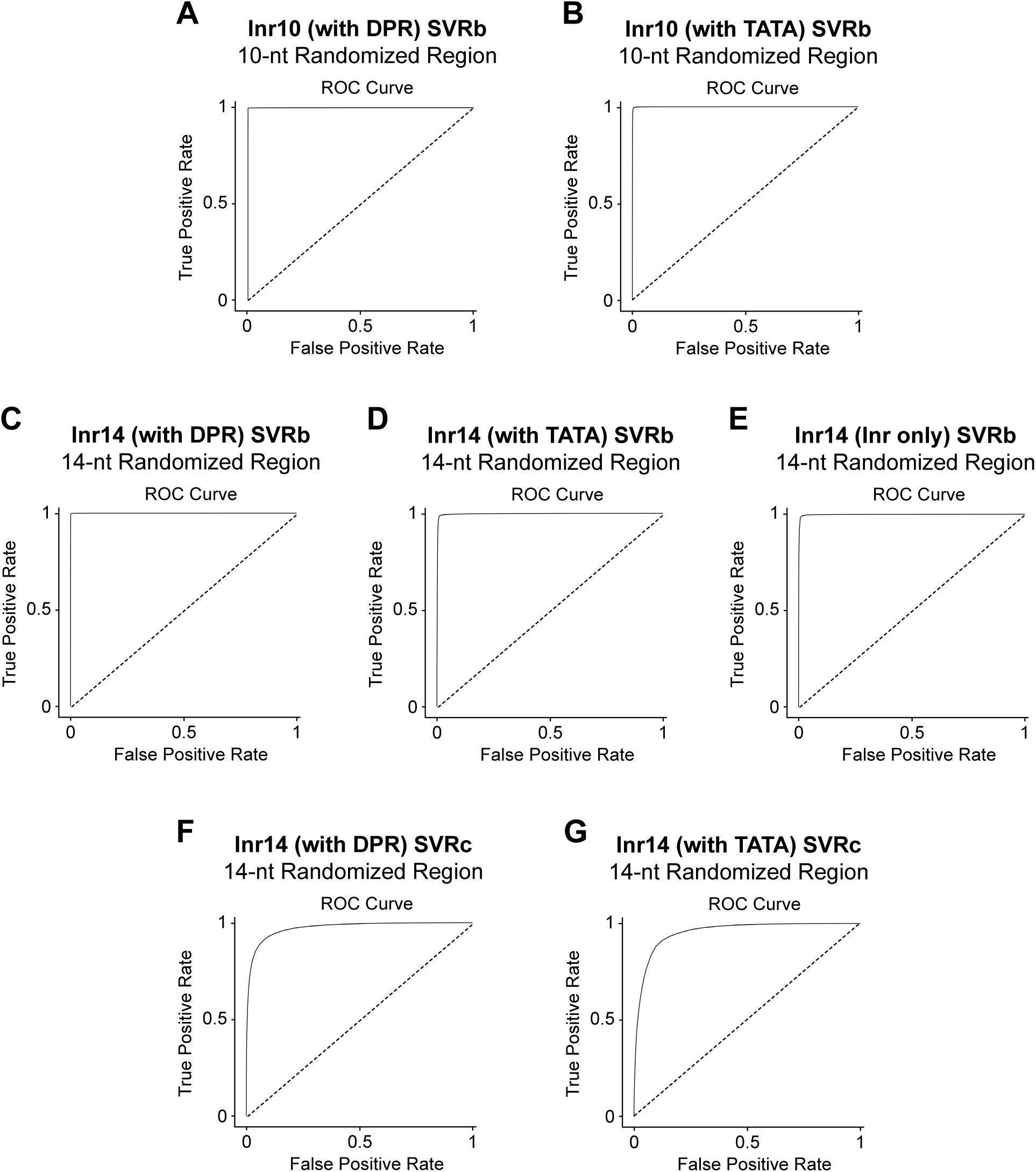
Receiver operating characteristic (ROC) curves of the Inr SVR models. The HARPE sequence variants used for the ROC curves are the same as in Supplemental Fig. S7. (*A–G*) ROC analysis of the SVR models described in Supplemental Fig. S7. Performance was computed by counting true positives (TP), true negatives (TN), false positives (FP), and false negatives (FN). True positive rate (or sensitivity or recall) [TP / (TP + FN)] is the proportion of transcriptionally active variants that are correctly predicted as positives. False positive rate [FP / (FP + TN)] is the probability for an inactive sequence to be incorrectly predicted as positive.

**Supplemental Figure S10.**
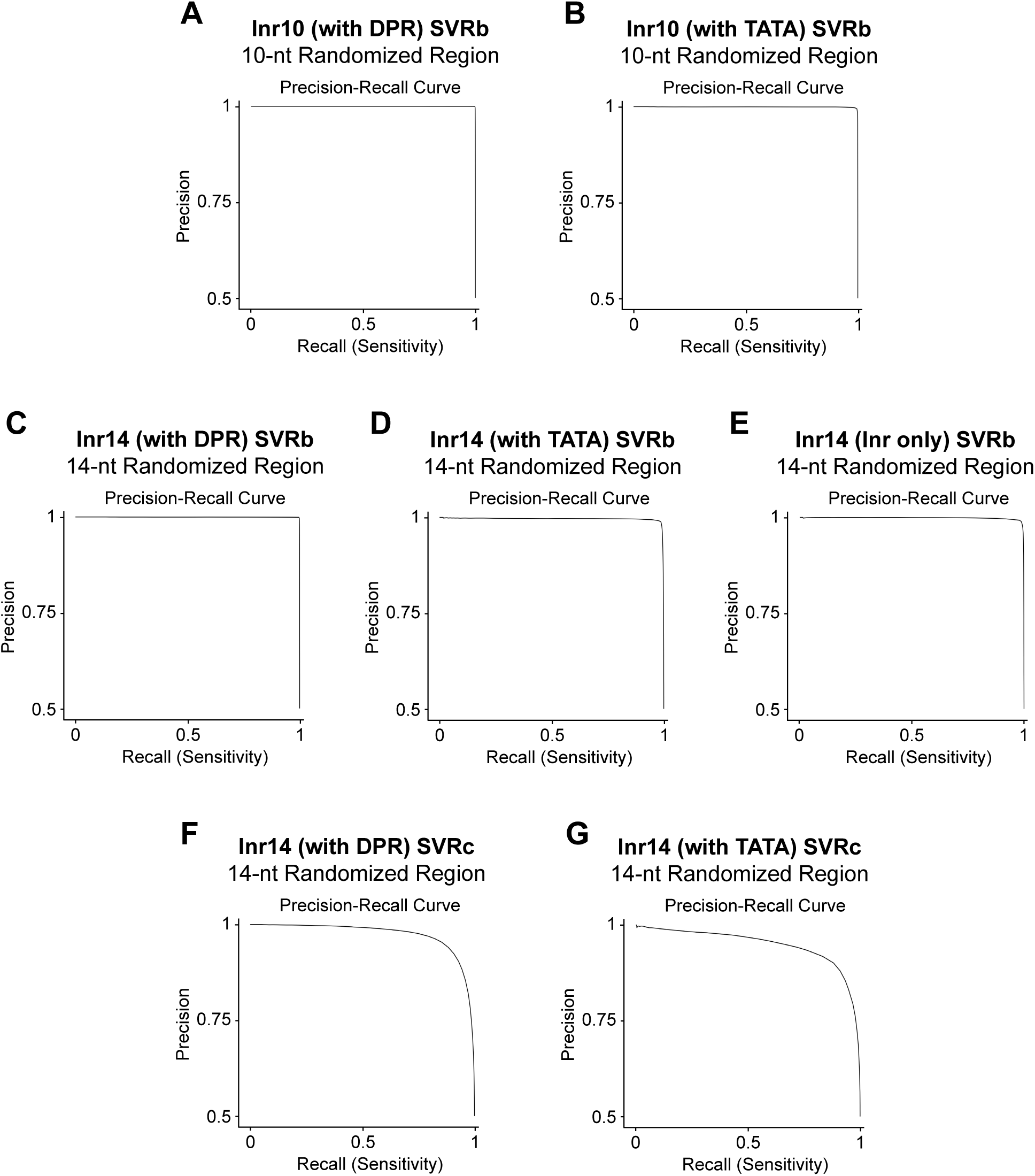
Precision-recall (PR) curves of the Inr SVR models. The HARPE sequence variants used for the PR curves are the same as in Supplemental Fig. S7. (*A–G*) Precision-recall curves of the SVR models described in Supplemental Fig. S7. Performance was computed by counting true positives (TP), true negatives (TN), false positives (FP), and false negatives (FN). Precision [TP / (TP + FP)] is the proportion of positive predictions that are correct. Recall (or sensitivity or true positive rate) [TP / (TP + FN)] is the proportion of transcriptionally active variants that are correctly predicted as positives.

**Supplemental Figure S11.**
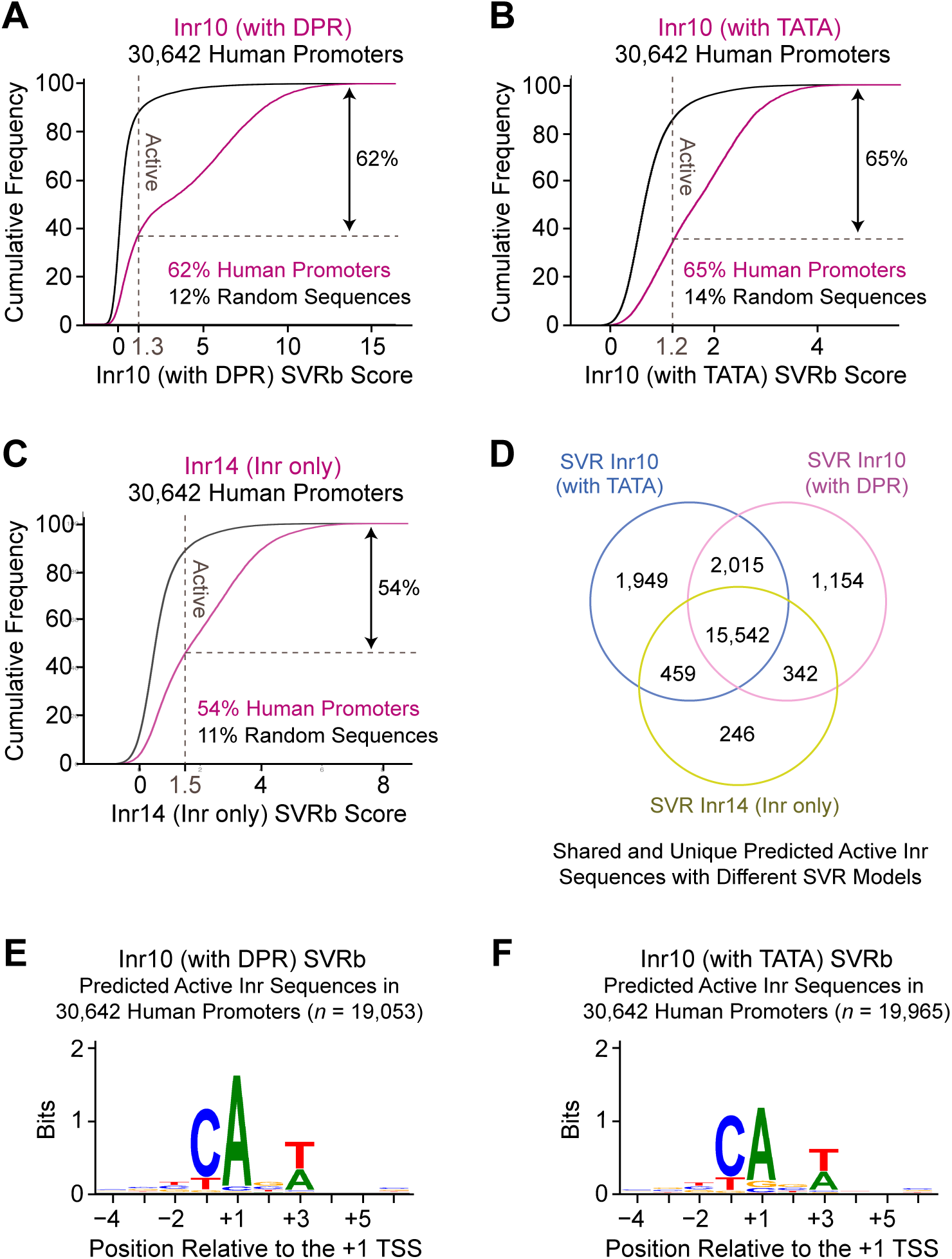
Approximately 60% of human promoters are predicted to contain an active Inr element. (*A–C*) Shown are the cumulative frequencies of Inr SVR scores of sequences in the Inr region (−4 to +6 or −5 to +9) of 30,642 natural human promoters. (*A*) With the Inr10 (with DPR) SVRb model, -62% of human promoters are predicted to have an active Inr (SVR 2’ 1.3), compared to only-12% of 100,000 random 10-nt sequences with the same overall G/C content (61%). (*B*) With the Inr10 (with TATA) SVRb model, -65% of human promoters are predicted to have an active Inr (SVR 2’ 1.2), compared to only -14% of 100,000 random 10-nt sequences with the same overall G/C content. (*C*) With the Inr14 (Inr only) SVRb model, -54% of human promoters are predicted to have an active Inr (SVR 2’ 1.5), compared to only -11% of 100,000 random 14-nt sequences with the same overall G/C content. (*D*) The majority of predicted active Inr sequences are shared across the three Inr SVR models. (*E,F*) Active Inr elements have a minimal consensus sequence of CA_+1_NW. Shown are sequence logos of the Inr region (−4 to +6) of 30,642 natural human promoters that are predicted to be active with the Inr10 (with DPR) SVRb model (*E*) or with the Inr10 (with TATA) SVRb model (*F*). TSSs were identified by using the GRO-cap method in human GM12878 cells.

**Supplemental Figure S12.**
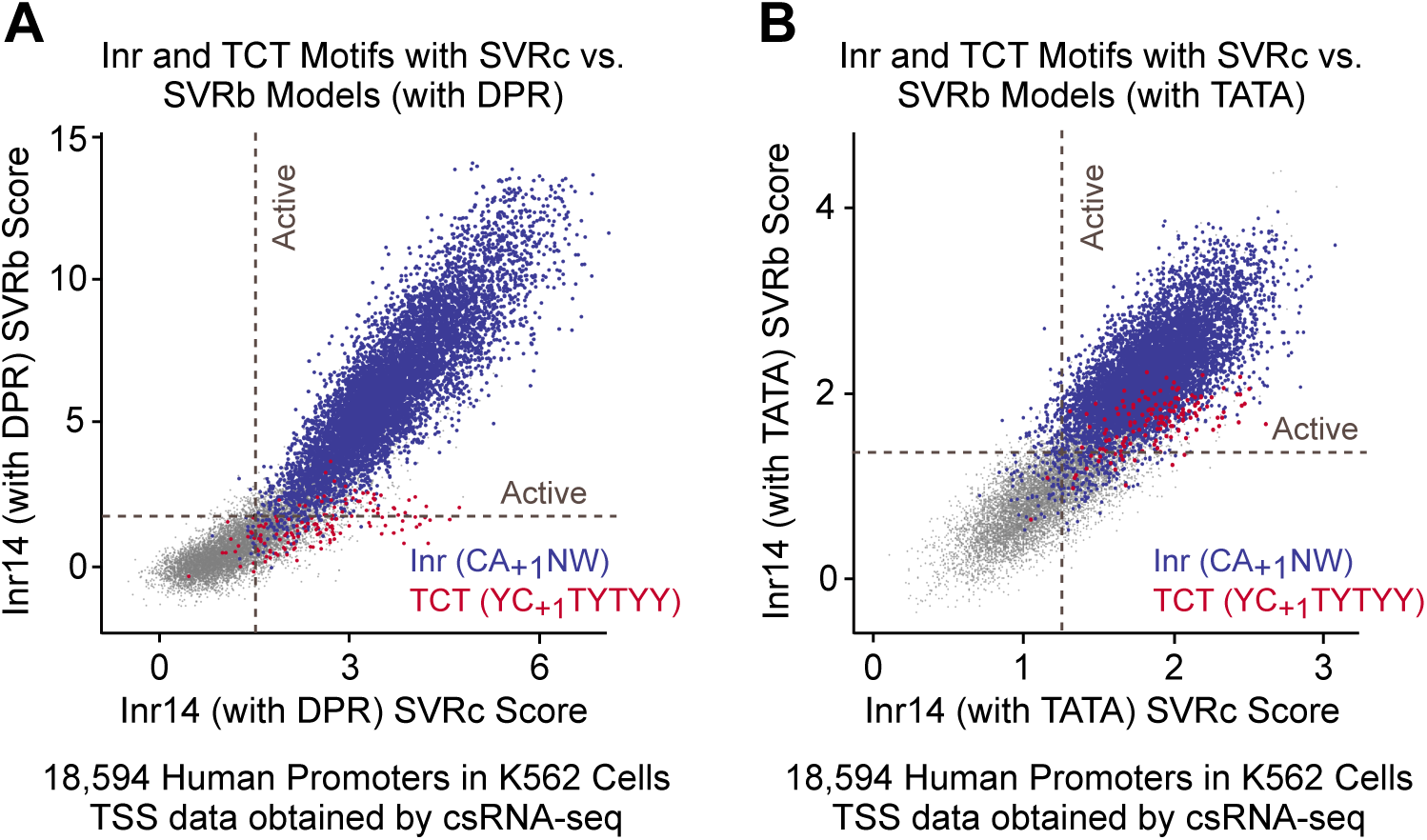
The TCT core promoter motif, which overlaps with the Inr motif, can be identified more distinctly with the cell-based Inr 14 SVRc models than with the biochemical Inr14 SVRb models. The Inr14 SVR models were generated with HARPE data obtained from promoters with a 14-nt randomized region (from −5 to +9 relative to the SCP1 +1 TSS). SVR scores are shown for 18,594 natural human promoters, with sequences matching the minimal Inr (CA_+1_NW) or TCT (YC_+1_TYTYY) motifs highlighted in blue or red, respectively. The red dots are plotted on top of the blue dots for visibility. (*A*) Comparison of Inr14 SVR models that were generated with a DPR-driven core promoter. (*B*) Comparison of Inr14 SVR models that were generated with a TATA-driven core promoter.

**Supplemental Figure S13.**
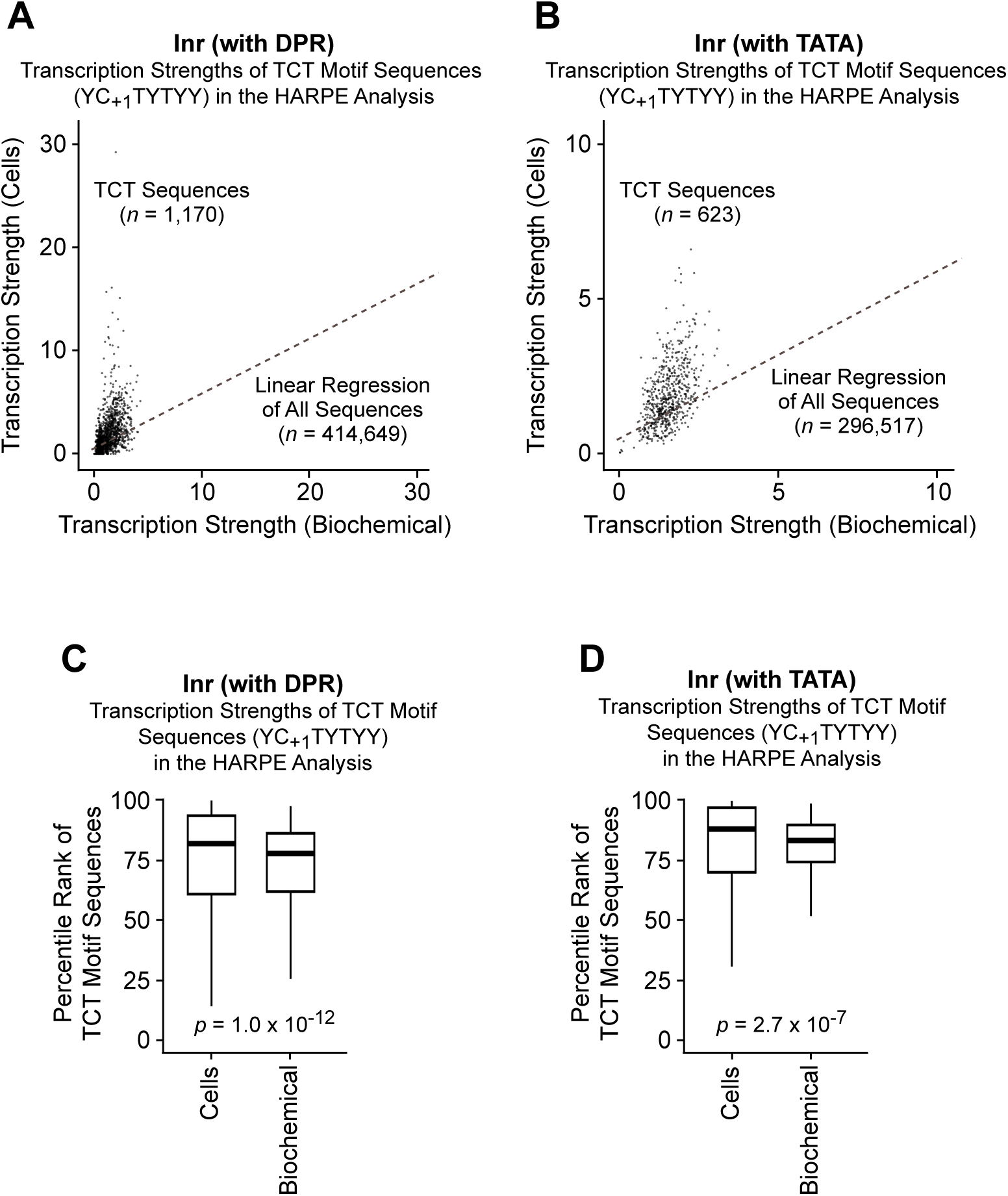
HARPE sequence variants that match the TCT motif (YC_+1_TYTYY) exhibit higher transcriptional activity in cells than in vitro. (*A,B*) Observed transcription strengths of TCT motif sequences in the cell-based versus biochemical HARPE experiments with a 14-nt randomized region (−5 to +9) in a DPR-driven (*A*) or TATA-driven (*B*) promoter. The dashed lines show the linear regression of all HARPE sequence variants that were detected in both the cell-based and biochemical experiments. (*C,D*) Percentile ranks of the observed transcription strengths of TCT motif sequences in the cell-based versus biochemical HARPE experiments with a 14-nt randomized region (−5 to +9) in a DPR-driven (*C*) or TATA-driven (*D*) promoter. The thick horizontal lines are the medians, and the lower and upper hinges are the first and third quartiles, respectively. Whiskers extend from the hinges to the largest or lowest value no further than 1.5 * IQR from the hinge. Data beyond the end of the whiskers (outlying points) are omitted from the box plots. The *p*-values were calculated with a Wilcoxon rank-sum test to compare the percentile distributions.

**Supplemental Figure S14.**
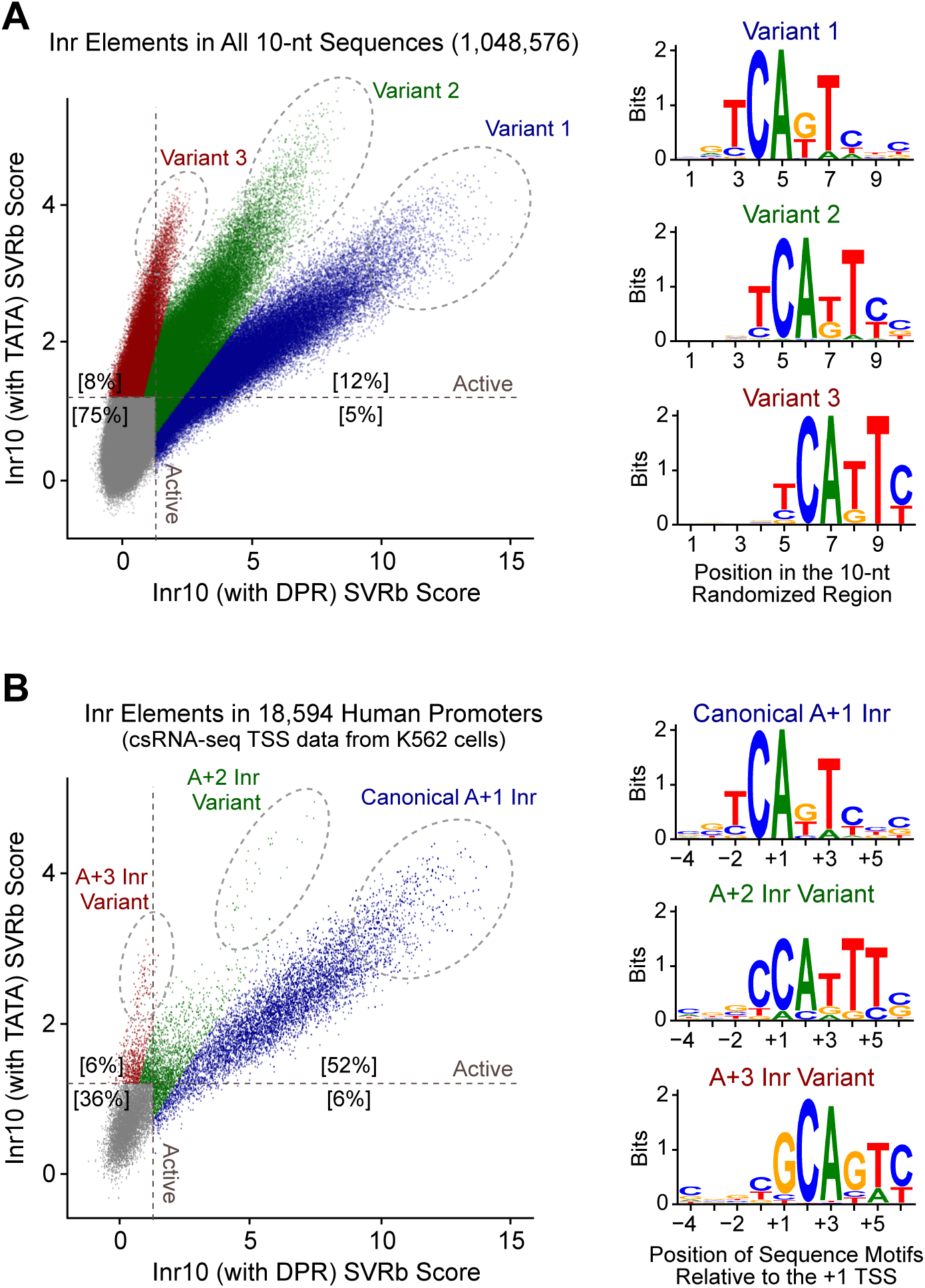
The positioning of the Inr sequence motif relative to the +1 TSS is different in DPR-driven promoters than in TATA-driven promoters. (*A*) Comparison of the predicted Inr activities of all possible 10-nt sequences (1,048,576) with the Inr (with DPR) SVR model versus the Inr (with TATA) SVR model reveals three distinct variants with different positioning of the core Inr CAKT motif within the 10-nt randomized region. The web logos are based on the sequences in the highlighted dashed ovals. (*B*) Comparison of the predicted Inr activities of 18,594 natural human promoters with the Inr (with DPR) SVR model versus Inr (with TATA) SVR model reveals the same three variants, which are labeled according to the position of the A nt in the core CAKT Inr motif relative to the observed +1 TSS.

**Supplemental Figure S15.**
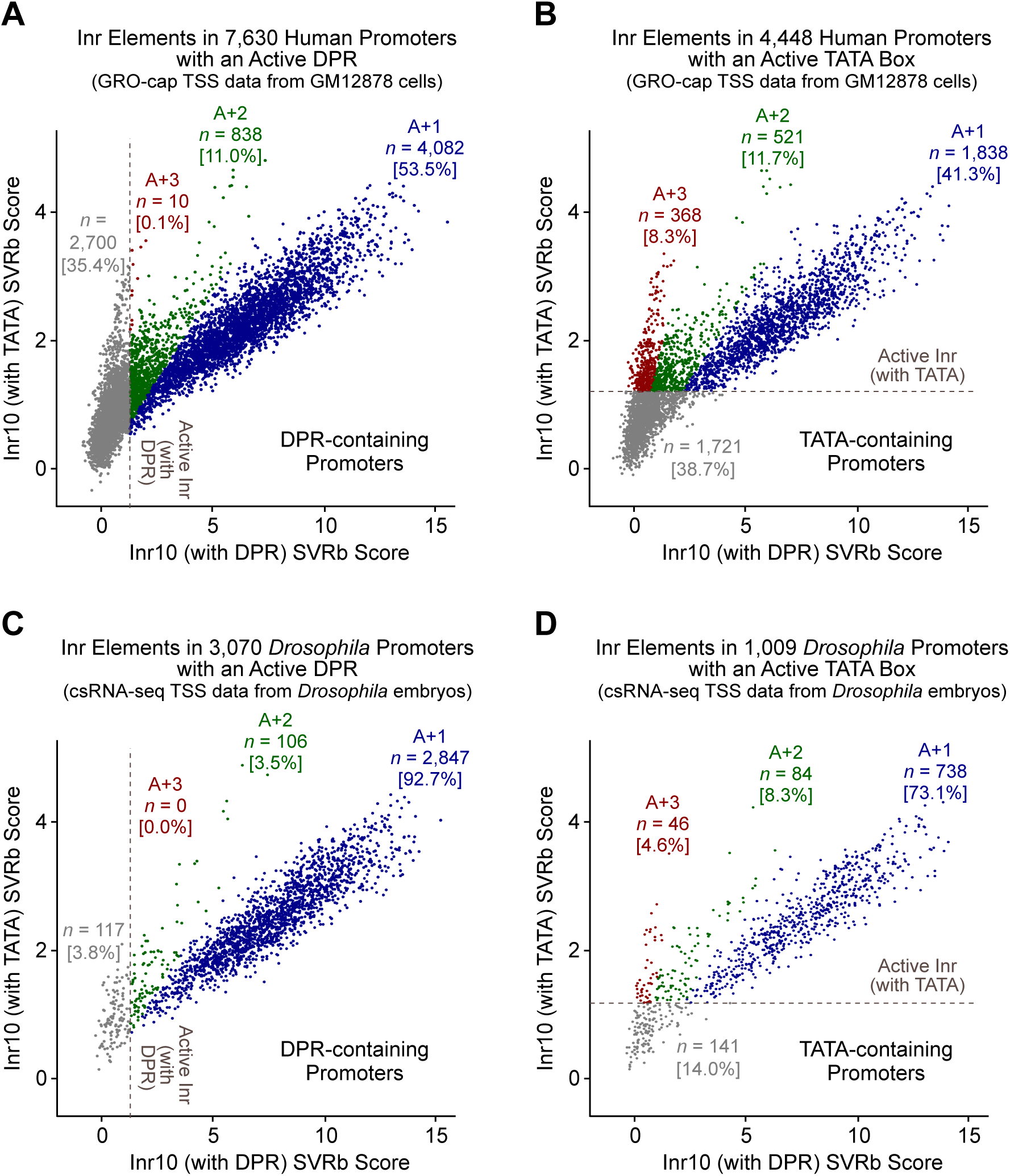
The positioning of the Inr sequence motif relative to the +1 TSS is different in natural DPR-containing promoters than in natural TATA-containing promoters. (*A,B*) Comparison of the predicted Inr activities of 7,630 DPR-containing (*A*) and 4,448 TATA-containing (*B*) human promoters with the Inr (with DPR) SVR model versus the Inr (with TATA) SVR models. (*C,D*) Comparison of the predicted Inr activities of 3,070 DPR-containing (*C*) and 1,009 TATA-containing (*D*) *Drosophila* promoters with the Inr (with DPR) SVR model versus the Inr (with TATA) SVR models. DPR-containing promoters were identified by using the SVR models of the human DPR (*A*) and the *Drosophila* DPR (*C*) (Vo ngoc et al. 2020, 2023). (*B,D*) TATA-containing promoters were identified by using the SVR model of the human TATA box (Vo ngoc et al. 2020).

**Supplemental Figure S16.**
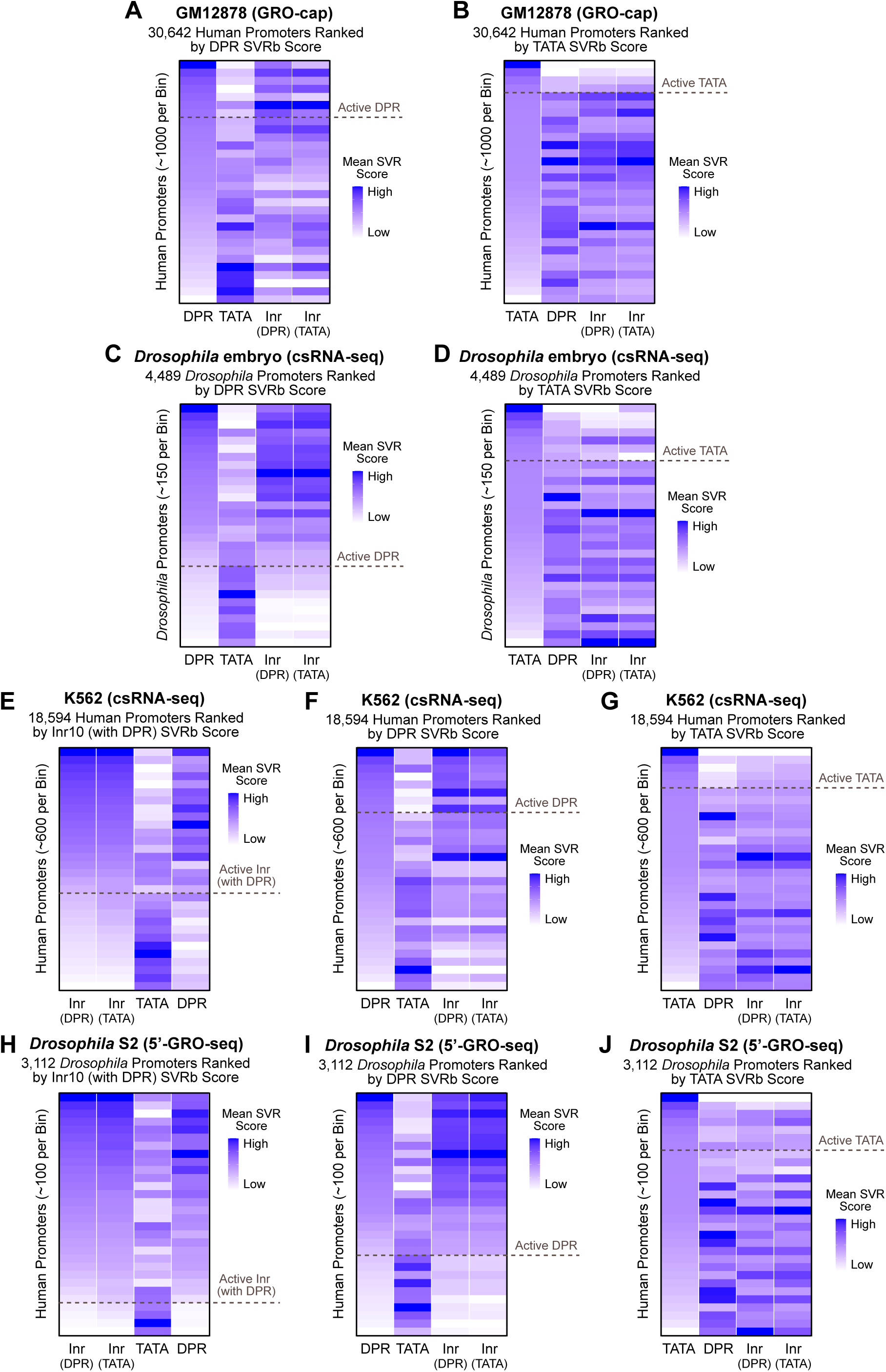
Predicted Inr activity positively correlates with predicted DPR activity and negatively correlates with predicted TATA box activity in humans as well as in *Drosophila*. Natural promoters were ranked according to the indicated SVR scores and divided into 30 bins of approximately equivalent size. For each bin, the mean SVR scores for human Inr (with DPR), human Inr (with TATA), human TATA box (Vo ngoc et al. 2020), and human or *Drosophila* DPR (Vo ngoc et al. 2020, 2023) were determined. (*A,B*) Predicted correlations in human GM12878 cells. TSSs were identifed by using the GRO-cap method. (*C,D*) Predicted correlations in *Drosophila* embryos. TSSs were identified by using the csRNA-seq method. (*E–G*) Predicted correlations in human K562 cells. TSSs were identified by using the csRNA-seq method. (*H–J*) Predicted correlations in *Drosophila* S2 cells. TSSs were identified by using the 5’-GRO-seq method.

**Supplemental Figure S17.**
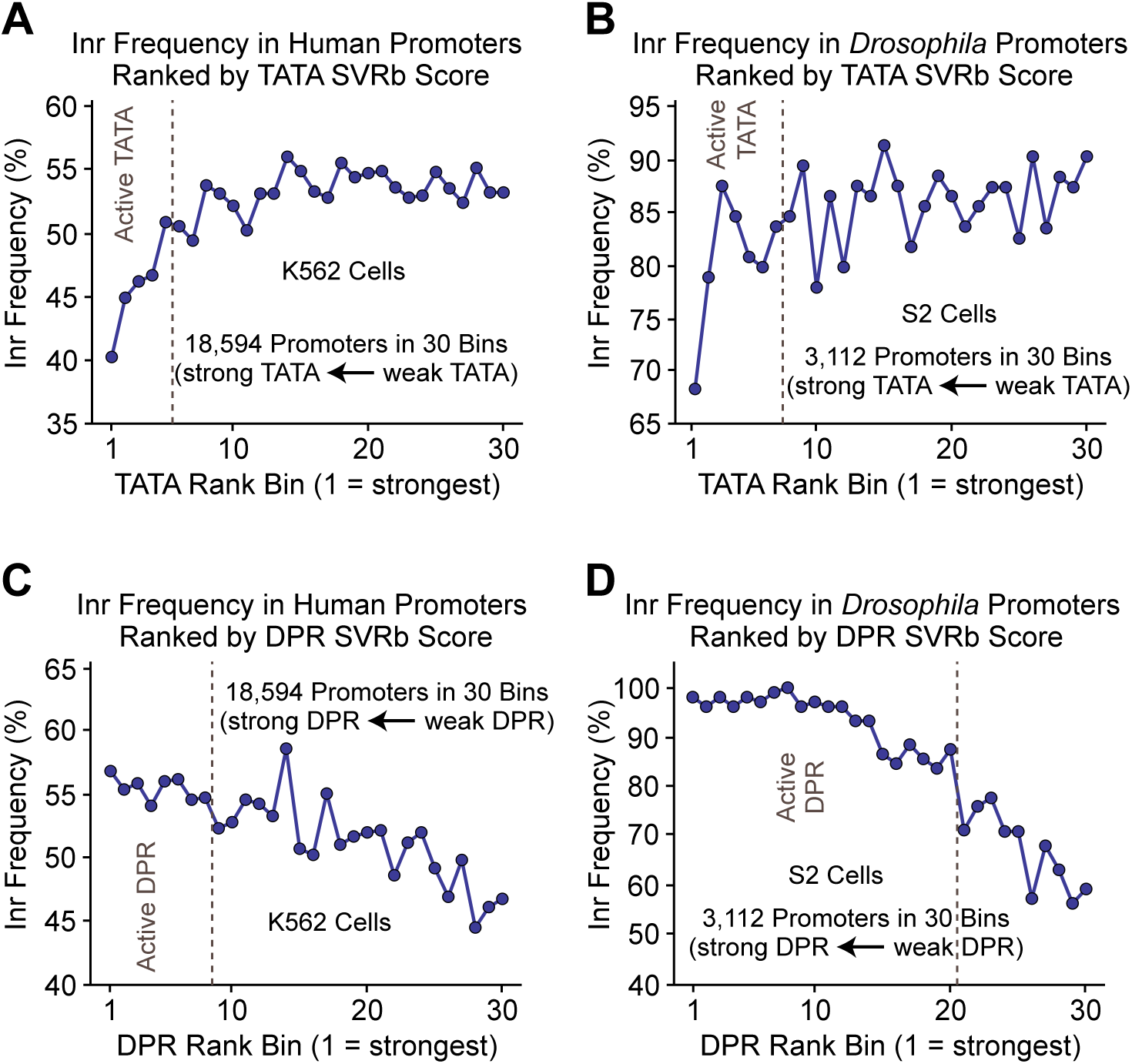
The predicted frequency of active Inr elements decreases with the predicted strength of TATA motifs and increases with the predicted strength of DPR motifs in human K562 cells as well as in *Drosophila* S2 cells. Shown are the predicted frequencies of active Inr elements as a function of the predicted strength of the TATA box (*A,B*) or DPR (*C,D*) in human (*A,C*) and *Drosophila* (*B,D*) promoters. Natural promoters were ranked according to their human TATA box scores (Vo ngoc et al. 2020) or human or *Drosophila* DPR scores (Vo ngoc et al. 2020, 2023) and divided into 30 bins of approximately equivalent size. Active Inr elements were defined to be those that are predicted to be active with both the Inr10 (with DPR) and Inr10 (with TATA) SVR models.

**Supplemental Figure S18.**
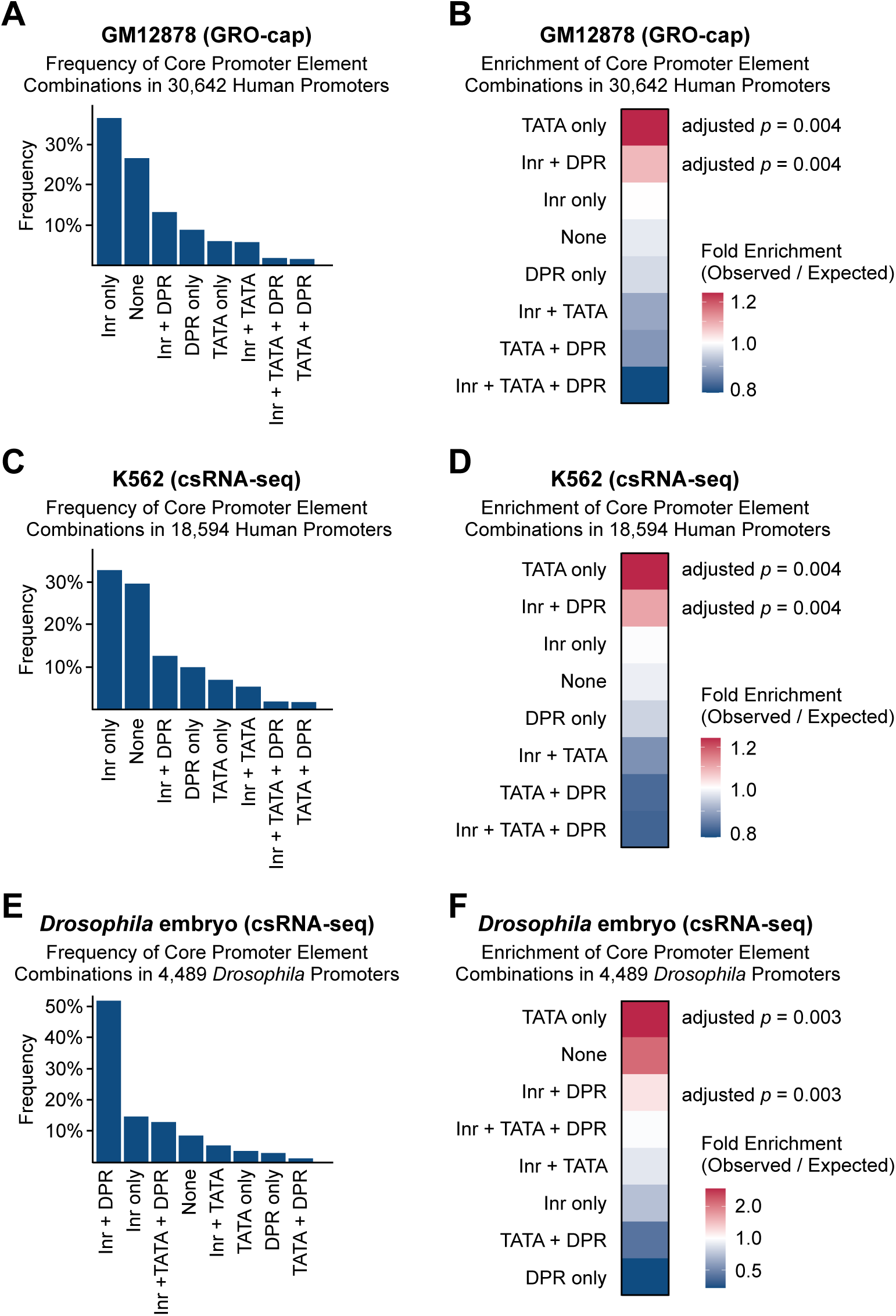
Predicted frequency and enrichment of core promoter element combinations in human and *Drosophila* promoters. The predicted frequency of each combination of elements in natural promoters was determined by using the SVRb models of the human Inr, human TATA box (Vo ngoc et al. 2020), and human or *Drosophila* DPR (Vo ngoc et al. 2020, 2023). Active Inr elements were defined to be those that are predicted to be active with both the Inr10 (with DPR) and Inr10 (with TATA) SVR models. The predicted enrichment of each combination of elements was determined by comparing its observed frequency in natural promoters to the expected frequency based on permutation testing (*n* = 1,000 permutations), where the binary activity value (active or inactive) for each element was shuffled between promoters. The *p*-values reflect the proportion of permutations with frequencies equal to or greater than observed (0 out of 1,000 for all values shown) and were adjusted for multiple comparisons by using the Benjamini-Hochberg correction. (*A,B*) Core promoter combinations based on TSSs that were identified by using the GRO-cap method in human GM12878 cells. (*C,D*) Core promoter combinations based on TSSs that were identified by using the csRNA-seq method in human K562 cells. (*E,F*) Core promoter combinations based on TSSs that were identified by using the csRNA-seq method in *Drosophila* embryos.

**Supplemental Figure S19.**
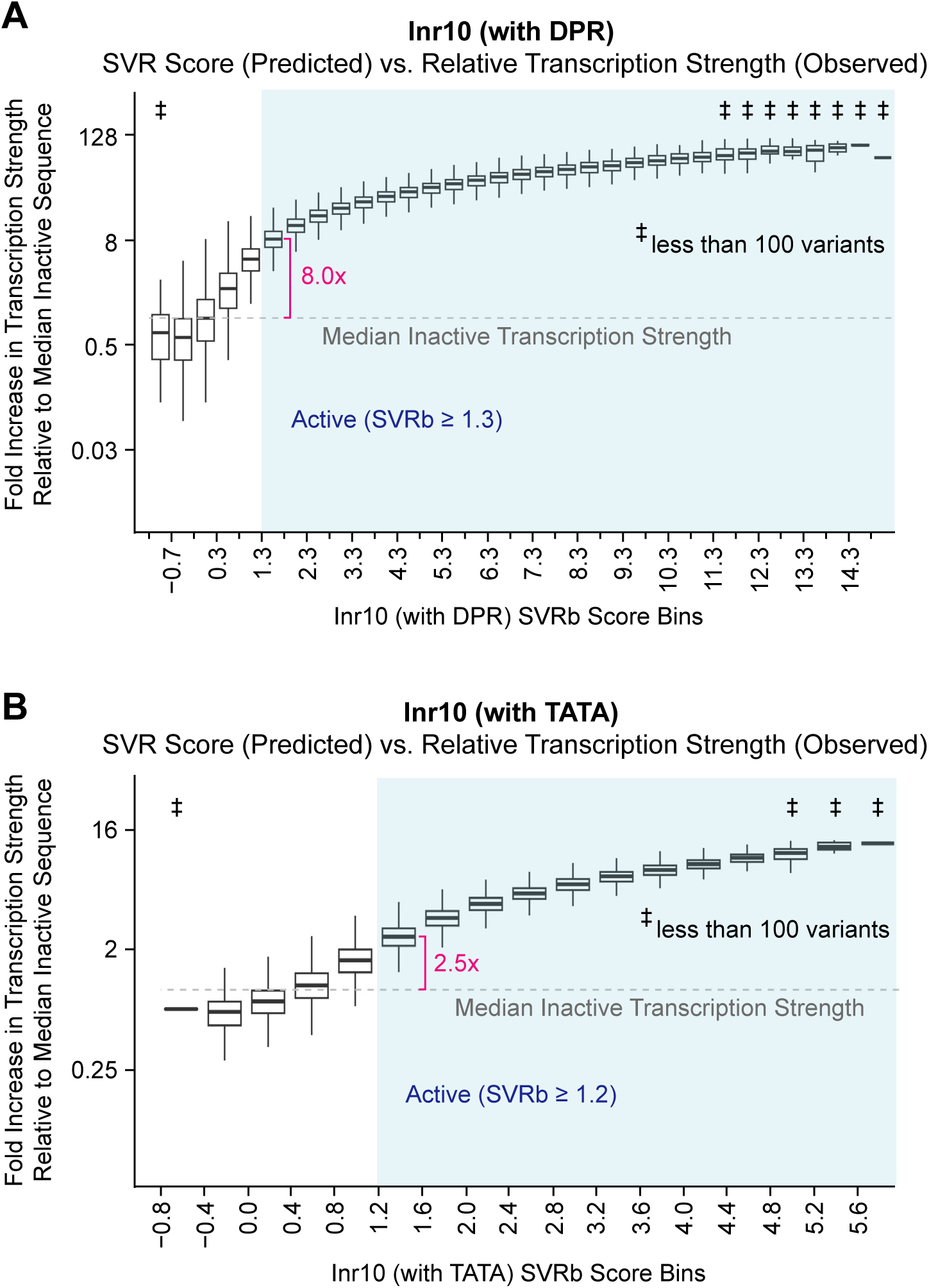
Inr sequence variants exhibit a broader range of transcriptional activity in DPR-driven promoters than in TATA-driven promoters. Shown are box-plot diagrams of the transcription strengths of all HARPE sequence variants placed into bins of the indicated SVR score ranges. (*A*) Sequence variants with Inr10 (with DPR) SVRb scores ≥ 1.3 are typically at least 8 times more active than an inactive sequence. (*B*) Sequence variants with Inr10 (with TATA) SVRb scores ≥1.2 are typically at least 2.5 times more active than an inactive sequence. In both diagrams, the thick horizontal lines are the medians, and the lower and upper hinges are the first and third quartiles, respectively. Whiskers extend from the hinges to the largest or lowest value no further than 1.5 * IQR from the hinge. Data beyond the end of the whiskers (outlying points) are omitted from the box-plots. Variants with transcription strength = 0 were removed to allow log-scale display of the diagram. The light blue shaded regions indicate sequence variants with SVR scores greater than or equal to the optimal SVR thresholds identified in Supplemental Fig. S7. The horizontal dashed grey line denotes the median transcription strength of inactive sequences.

**Supplemental Figure S20.**
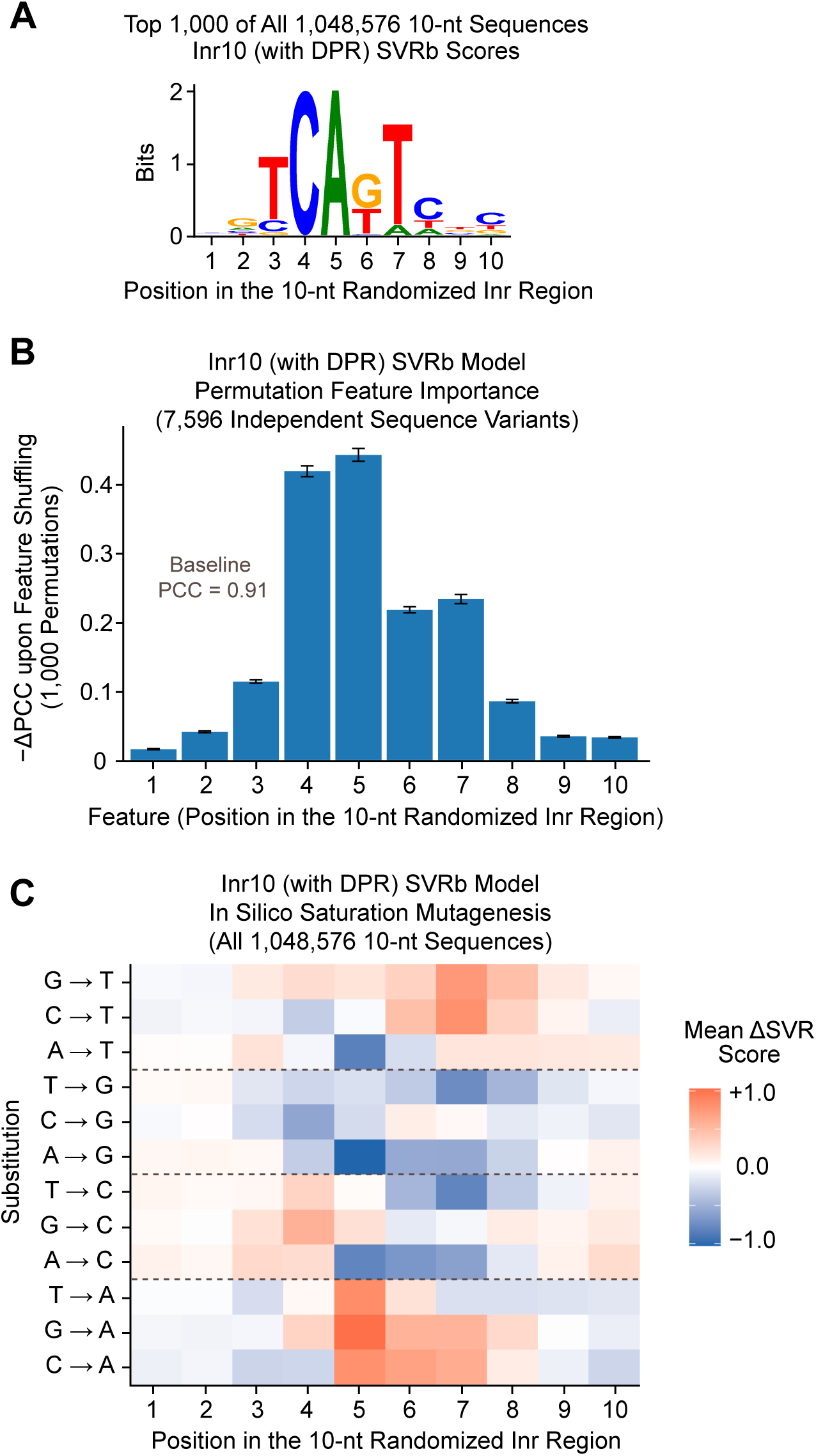
Model interpretation analyses reveal key sequence features learned by the Inr10 (with DPR) SVRb model. (*A*) SVR analysis of all possible 10-nt sequences reveals the predicted optimal Inr motif. Shown is the web logo of the predicted top 1,000 out of all 1,048,576 10-nt sequences. (*B*) Permutation feature importance analysis reveals that positions 4 to 7, which correspond to the core Inr CAKT motif, are most important for SVR prediction of transcriptional activity. Each feature (position in the 10-nt randomized Inr region) was shuffled between the 7,596 independent test sequences shown in Fig. 1D (left), and the resulting decrease in model performance [as indicated by the change in Pearson’s correlation coefficient (PCC) between the observed transcription strengths and predicted SVR scores] was measured. The baseline model performance (PCC = 0.91) is shown in Fig. 1D (left). (*C*) In silico saturation mutagenesis reveals position- and nucleotide-specific effects on predicted transcriptional activity. Each position in the 10-nt randomized Inr region was systematically mutated across all possible sequences to each of the three alternative nucleotides. Shown is the mean change in SVR score for each substitution.

**Supplemental Figure S21.**
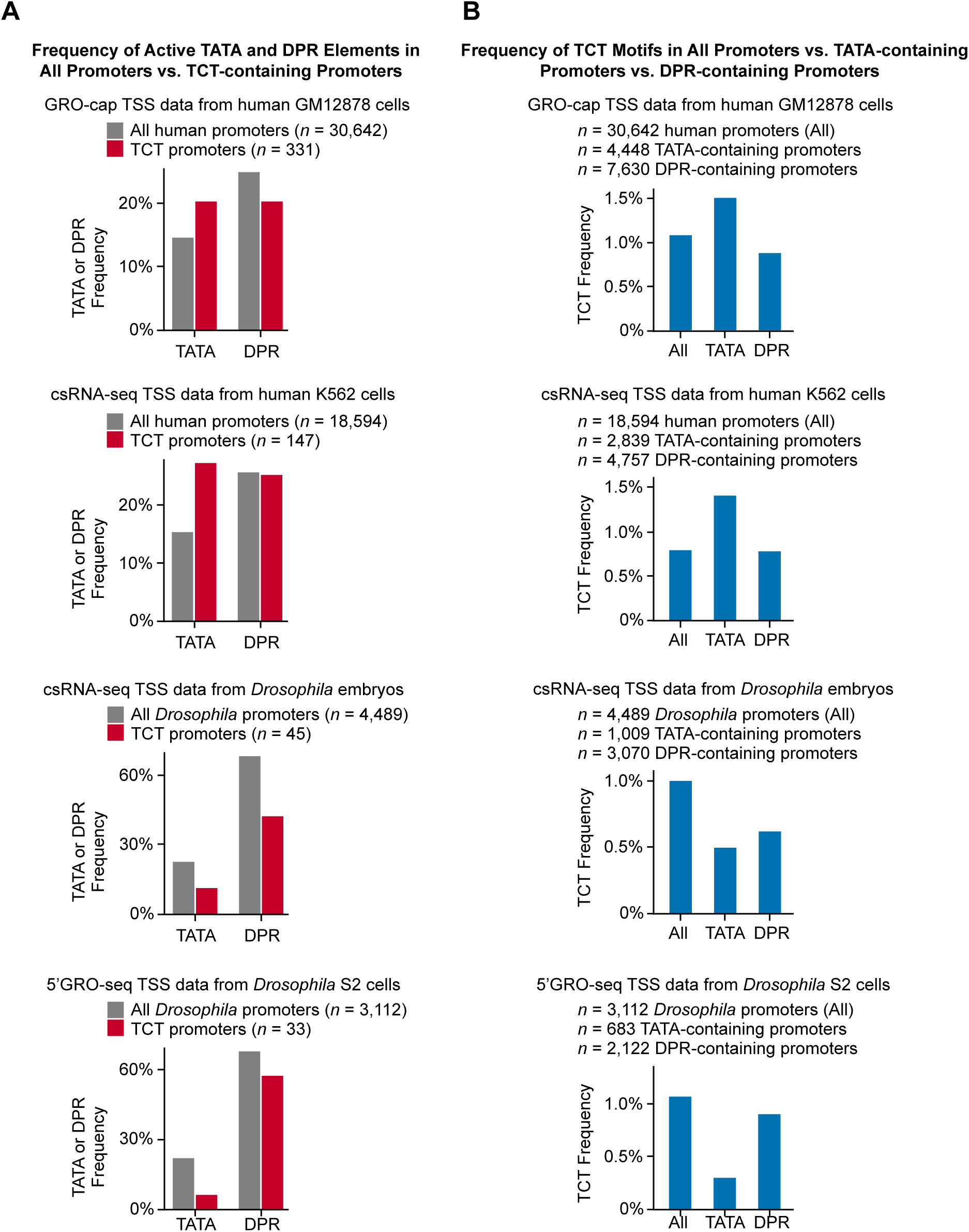
The TCT motif functions differently in humans than in *Drosophila*. (*A*) The TATA box is overrepresented in human TCT promoters and is underrrepresented in *Drosophila* TCT promoters. The predicted frequencies of active TATA and DPR elements in all identified promoters versus only TCT-containing promoters are shown. (*B*) The TCT motif is enriched in human TATA-containing promoters and is depleted in *Drosophila* TATA-containing promoters. The frequencies of TCT motifs in all identified promoters (All), TATA-containing promoters, and DPR-containing promoters are shown. The predicted activities of the TATA and DPR elements were determined by using SVR models of the human TATA box (Vo ngoc et al. 2020) and the human and *Drosophila* DPR (Vo ngoc et al. 2020, 2023). TCT motifs were identified by a match to the TCT consensus sequence (YC_+1_TYTYY in humans and YYC_+1_TTTYY in *Drosophila*).

**Supplemental Table 1.**

| Promoter Construct | HARPE Experiment |  | Reproducibility (PCC)<br>(transcription strength; rep1 vs rep 2) |
| --- | --- | --- | --- |
|  | Randomized Region | Biochemical or Cells |  |
| Inr10 (with DPR) | 10-nt | Biochemical | 0.94 |
| Inr10 (with TATA) | 10-nt | Biochemical | 0.91 |
| Inr14 (with DPR) | 14-nt | Biochemical | 0.96 |
| Inr14 (with TATA) | 14-nt | Biochemical | 0.89 |
| Inr14 (Inr only) | 14-nt | Biochemical | 0.71, 0.60, 0.62 * |
| Inr14 (with DPR) | 14-nt | Cells | 0.55 |
| Inr14 (with TATA) | 14-nt | Cells | 0.63 |

\*Due to low transcription levels with this promoter, we performed 3 replicates. The three PCC values are for rep1 vs. rep2, rep2 vs. rep3, and rep1 vs. rep3.

**Supplemental Table 2.**

| GM12878 cells (GRO-cap TSS data) |  |  |  |  |  |  |
| --- | --- | --- | --- | --- | --- | --- |
| Human Promoter Types | % Active Inr10 (with DPR) | % Active Inr10 (with TATA) | % Active Inr (either)* | % Active Inr (both)** | % Active TATA | % Active DPR |
| All | 62.2 | 65.2 | 70 | 57.3 | 14.5 | 24.9 |
| Active Inr10 (with DPR) | [100] | 92.1 | [100] | 92.1 | 12.7 | 25.9 |
| Active Inr10 (with TATA) | 87.9 | [100] | [100] | 87.9 | 13.7 | 25.4 |
| Active Inr (either)* | 88.8 | 93 | [100] | 81.8 | 13.5 | 25.4 |
| Active Inr (both)** | [100] | [100] | [100] | [100] | 12.7 | 25.9 |
| Inactive Inr (both)*** | [0] | [0] | [0] | [0] | 16.8 | 23.7 |
| Active TATA | 54.1 | 61.3 | 65.3 | 50.1 | [100] | 20.8 |
| Inactive TATA | 63.5 | 65.8 | 70.8 | 58.5 | [0] | 25.6 |
| Active DPR | 64.6 | 66.4 | 71.5 | 59.5 | 12.1 | [100] |
| Inactive DPR | 61.4 | 64.7 | 69.5 | 56.6 | 15.3 | [0] |

\*Active with the Inr10 (with TATA) SVR model and/or the Inr10 (with DPR) SVR model
\*\*Active with both the Inr10 (with TATA) SVR model and the Inr10 (with DPR) SVR model
\*\*\*Inactive with both the Inr10 (with TATA) SVR model and the Inr10 (with DPR) SVR model

| K562 cells (csRNA-seq TSS data) |  |  |  |  |  |  |
| --- | --- | --- | --- | --- | --- | --- |
| Human Promoter Types | % Active Inr10 (with DPR) | % Active Inr10 (with TATA) | % Active Inr (either)* | % Active Inr (both)** | % Active TATA | % Active DPR |
| All | 58.4 | 58.1 | 64.3 | 52.2 | 15.3 | 25.6 |
| Active Inr10 (with DPR) | [100] | 89.4 | [100] | 89.4 | 13.4 | 27.1 |
| Active Inr10 (with TATA) | 89.8 | [100] | [100] | 89.8 | 13.9 | 26.7 |
| Active Inr (either)* | 90.8 | 90.4 | [100] | 81.1 | 14 | 26.7 |
| Active Inr (both)** | [100] | [100] | [100] | [100] | 13.2 | 27.2 |
| Inactive Inr (both)*** | [0] | [0] | [0] | [0] | 17.5 | 23.6 |
| Active TATA | 51.2 | 53.0 | 59.0 | 45.1 | [100] | 21.3 |
| Inactive TATA | 59.7 | 59.1 | 65.3 | 53.5 | [0] | 26.4 |
| Active DPR | 61.8 | 60.7 | 67.1 | 55.5 | 12.7 | [100] |
| Inactive DPR | 57.2 | 57.3 | 63.4 | 51.1 | 16.2 | [0] |

\*Active with the Inr10 (with TATA) SVR model and/or the Inr10 (with DPR) SVR model
\*\*Active with both the Inr10 (with TATA) SVR model and the Inr10 (with DPR) SVR model
\*\*\*Inactive with both the Inr10 (with TATA) SVR model and the Inr10 (with DPR) SVR model

| *Drosophila* embryos (csRNA-seq TSS data)**** |  |  |  |  |  |  |
| --- | --- | --- | --- | --- | --- | --- |
| *Drosophila* Promoter Types | % Active Inr10 (with DPR) | % Active Inr10 (with TATA) | % Active Inr (either)* | % Active Inr (both)** | % Active TATA | % Active DPR |
| All | 86.6 | 87.9 | 90.2 | 84.3 | 22.5 | 68.4 |
| Active Inr10 (with DPR) | [100] | 97.2 | [100] | 97.2 | 21.3 | 75.9 |
| Active Inr10 (with TATA) | 95.9 | [100] | [100] | 95.9 | 22 | 74.1 |
| Active Inr (either)* | 96 | 97.4 | [100] | 93.4 | 22 | 73.6 |
| Active Inr (both)** | [100] | [100] | [100] | [100] | 21.3 | 76.6 |
| Inactive Inr (both)*** | [0] | [0] | [0] | [0] | 26.7 | 20.5 |
| Active TATA | 82.3 | 86 | 88.4 | 79.9 | [100] | 61.3 |
| Inactive TATA | 87.9 | 88.4 | 90.8 | 85.5 | [0] | 70.4 |
| Active DPR | 96.2 | 95.2 | 97.1 | 94.3 | 20.2 | [100] |
| Inactive DPR | 66.0 | 72.0 | 75.5 | 62.4 | 27.5 | [0] |

\*Active with the human Inr10 (with TATA) SVR model and/or the human Inr10 (with DPR) SVR model
\*\*Active with both the human Inr10 (with TATA) SVR model and the human Inr10 (with DPR) SVR model
\*\*\*Inactive with both the Inr10 (with TATA) SVR model and the Inr10 (with DPR) SVR model
\*\*\*\*Using the *Drosophila* DPR SVR model and the human Inr10 & TATA SVR models

| S2 cells (5'GRO-seq TSS data)**** |  |  |  |  |  |  |
| --- | --- | --- | --- | --- | --- | --- |
| *Drosophila* Promoter Types | % Active Inr10 (with DPR) | % Active Inr10 (with TATA) | % Active Inr (either)* | % Active Inr (both)** | % Active TATA | % Active DPR |
| All | 86.7 | 87.2 | 89.1 | 84.8 | 21.9 | 68.2 |
| Active Inr10 (with DPR) | [100] | 97.8 | [100] | 97.8 | 20.9 | 74.6 |
| Active Inr10 (with TATA) | 97.9 | [100] | [100] | 97.9 | 21.1 | 74.0 |
| Active Inr (either)* | 97.3 | 97.8 | [100] | 95.1 | 21.3 | 73.2 |
| Active Inr (both)** | [100] | [100] | [100] | [100] | 20.8 | 74.8 |
| Inactive Inr (both)*** | [0] | [0] | [0] | [0] | 27.5 | 26.9 |
| Active TATA | 82.6 | 84.3 | 86.4 | 80.5 | [100] | 63.1 |
| Inactive TATA | 87.9 | 88.0 | 89.9 | 86.0 | [0] | 69.6 |
| Active DPR | 94.8 | 93.9 | 95.7 | 93.0 | 20.3 | [100] |
| Inactive DPR | 69.3 | 72.9 | 75.1 | 67.2 | 25.5 | [0] |

\*Active with the human Inr10 (with TATA) SVR model and/or the human Inr10 (with DPR) SVR model
\*\*Active with both the human Inr10 (with TATA) SVR model and the human Inr10 (with DPR) SVR model
\*\*\*Inactive with both the Inr10 (with TATA) SVR model and the Inr10 (with DPR) SVR model
\*\*\*\*Using the *Drosophila* DPR SVR model and the human Inr10 & TATA SVR models

**Supplemental Table 3A.**
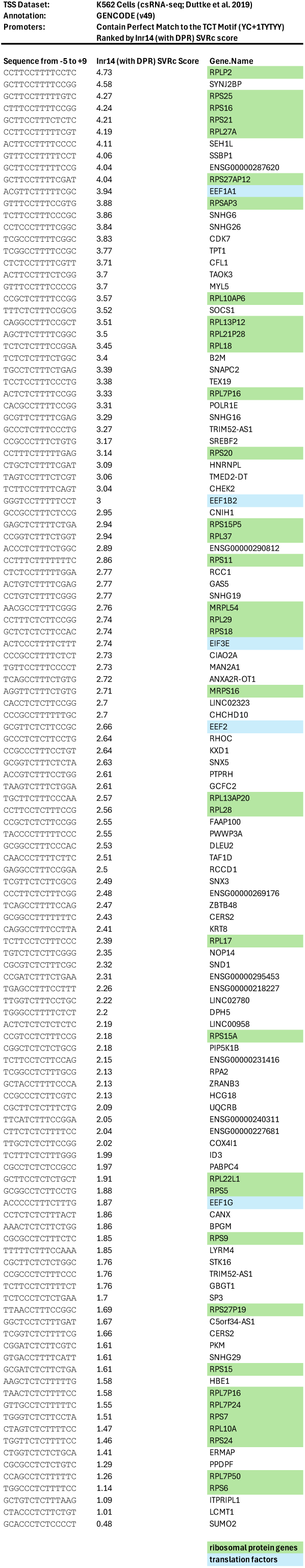

**Supplemental Table 3B.**
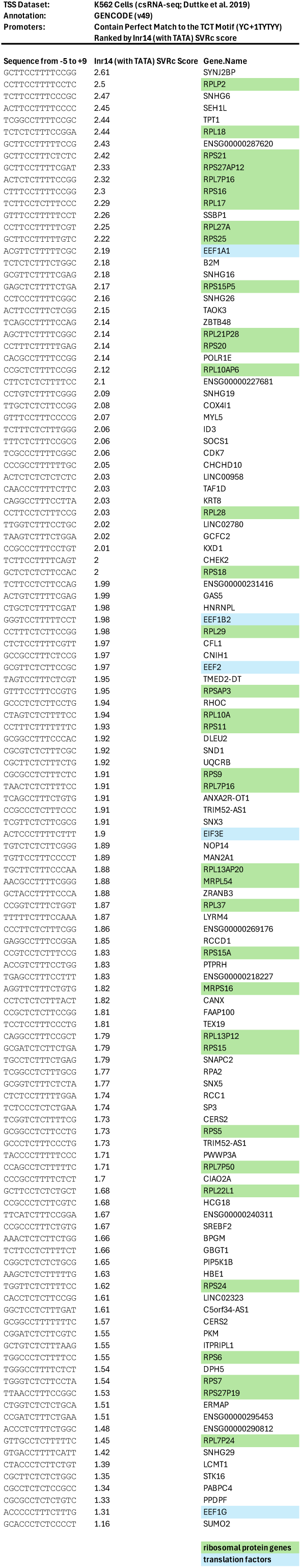

**Supplemental Table 4A.**

| ID | Description | GeneRatio | BgRatio | p.adjust |
| --- | --- | --- | --- | --- |
| GO:0010628 | positive regulation of gene expression | 68/498 | 491/6988 | 0.00014 |
| GO:0034097 | response to cytokine | 58/498 | 406/6988 | 0.00027 |
| GO:0071345 | cellular response to cytokine stimulus | 48/498 | 361/6988 | 0.01707 |
| GO:0006338 | chromatin remodeling | 47/498 | 358/6988 | 0.02115 |
| GO:0070848 | response to growth factor | 35/498 | 240/6988 | 0.02115 |
| GO:0000165 | MAPK cascade | 40/498 | 291/6988 | 0.02115 |
| GO:0030857 | negative regulation of epithelial cell differentiation | 6/498 | 11/6988 | 0.02132 |
| GO:0043408 | regulation of MAPK cascade | 34/498 | 237/6988 | 0.02473 |
| GO:0071363 | cellular response to growth factor stimulus | 33/498 | 228/6988 | 0.02473 |
| GO:0001656 | metanephros development | 8/498 | 22/6988 | 0.02721 |
| GO:0048871 | multicellular organismal-level homeostasis | 43/498 | 334/6988 | 0.02721 |
| GO:0043405 | regulation of MAP kinase activity | 13/498 | 55/6988 | 0.02721 |
| GO:0060993 | kidney morphogenesis | 8/498 | 23/6988 | 0.02969 |
| GO:0006979 | response to oxidative stress | 29/498 | 196/6988 | 0.02969 |
| GO:0072028 | nephron morphogenesis | 7/498 | 18/6988 | 0.03083 |
| GO:0072088 | nephron epithelium morphogenesis | 7/498 | 18/6988 | 0.03083 |
| GO:0043410 | positive regulation of MAPK cascade | 24/498 | 155/6988 | 0.04242 |
| GO:0045648 | positive regulation of erythrocyte differentiation | 6/498 | 14/6988 | 0.04242 |
| GO:0097190 | apoptotic signaling pathway | 40/498 | 317/6988 | 0.04242 |
| GO:0034612 | response to tumor necrosis factor | 19/498 | 111/6988 | 0.04242 |
| GO:0038034 | signal transduction in absence of ligand | 9/498 | 32/6988 | 0.04242 |
| GO:0097192 | extrinsic apoptotic signaling pathway in absence of ligand | 9/498 | 32/6988 | 0.04242 |
| GO:0006334 | nucleosome assembly | 11/498 | 46/6988 | 0.04242 |
| GO:0000302 | response to reactive oxygen species | 17/498 | 94/6988 | 0.04242 |
| GO:0009612 | response to mechanical stimulus | 14/498 | 70/6988 | 0.04641 |
| GeneRatio | fraction of input genes that are annotated to each GO term |  |  |  |
| BgRatio | fraction of background ("universe") genes that are annotated to each GO term |  |  |  |
| p.adjust | significance after Benjamini-Hochberg correction for multiple comparisons |  |  |  |

**Supplemental Table 4B.**

| ID | Description | setSize | NES | p.adjust |
| --- | --- | --- | --- | --- |
| GO:0006334 | nucleosome assembly | 46/6988 | 1.660116244 | 0.00008 |
| GO:0034728 | nucleosome organization | 60/6988 | 1.621213769 | 0.00008 |
| GO:0006338 | chromatin remodeling | 358/6988 | 1.328898765 | 0.00008 |
| GO:0065004 | protein-DNA complex assembly | 121/6988 | 1.465126199 | 0.00018 |
| GO:0048871 | multicellular organismal-level homeostasis | 334/6988 | 1.313360986 | 0.00051 |
| GO:1903706 | regulation of hemopoiesis | 197/6988 | 1.374269269 | 0.00058 |
| GO:0060284 | regulation of cell development | 333/6988 | 1.31177516 | 0.00058 |
| GO:0006325 | chromatin organization | 443/6988 | 1.276592398 | 0.00058 |
| GO:0000122 | negative regulation of transcription by RNA polymerase II | 415/6988 | 1.275706262 | 0.00058 |
| GO:0071363 | cellular response to growth factor stimulus | 228/6988 | 1.350200719 | 0.00072 |
| GO:0071345 | cellular response to cytokine stimulus | 361/6988 | 1.291639409 | 0.00084 |
| GO:0060429 | epithelium development | 382/6988 | 1.282729569 | 0.00084 |
| GO:0034097 | response to cytokine | 406/6988 | 1.278057771 | 0.00084 |
| GO:0009628 | response to abiotic stimulus | 474/6988 | 1.263211652 | 0.00084 |
| GO:0007155 | cell adhesion | 479/6988 | 1.256588566 | 0.00089 |
| GO:0010628 | positive regulation of gene expression | 491/6988 | 1.254221993 | 0.00117 |
| GO:0045910 | negative regulation of DNA recombination | 32/6988 | 1.648419289 | 0.00118 |
| GO:0040011 | locomotion | 405/6988 | 1.273776095 | 0.00123 |
| GO:0070848 | response to growth factor | 240/6988 | 1.332514242 | 0.00124 |
| GO:0001944 | vasculature development | 244/6988 | 1.314518151 | 0.00158 |
| GO:0051093 | negative regulation of developmental process | 323/6988 | 1.287451761 | 0.00158 |
| GO:0030261 | chromosome condensation | 28/6988 | 1.660573869 | 0.00162 |
| GO:1902105 | regulation of leukocyte differentiation | 158/6988 | 1.379658267 | 0.00162 |
| GO:0048514 | blood vessel morphogenesis | 198/6988 | 1.333561384 | 0.00162 |
| GO:0008285 | negative regulation of cell population proliferation | 267/6988 | 1.308574753 | 0.00162 |
| setSize | fraction of genes from the ranked list that are annotated to each GO term |  |  |  |
| NES | (normalized enrichment score) measure of the strength/direction of a GO term's distribution in the ranked list, with positive values indicating clustering towards the top |  |  |  |
| p.adjust | significance after Benjamini–Hochberg correction for multiple comparisons |  |  |  |

### Supplemental Materials and Methods

#### HARPE library generation

The Inr HARPE libraries were generated as described in Vo ngoc et al. (2020), except that some libraries were prepared by electroporation with MegaX DH10B T1R Electrocomp Cells (Thermo Fisher Scientific) instead of DH5G CloneCatcher Gold Electrocompetent Cells (Genlantis). The HARPE vector and SCP1-based promoter backbones, which include constructs with mutations in the DPR and/or the TATA box, are identical to those used previously (Vo ngoc et al. 2020, 2023). For HARPE analysis of the Inr, the 14-nt region from −5 to +9 and the 10-nt region from −4 to +6 (relative to the SCP1 +1 TSS) were randomized to generate ∼500,000 Inr sequence variants within SCP1 promoter backbones containing a DPR (mutTATA), a TATA box (mutDPR), or neither element (mutTATA, mutDPR). With the 10-nt libraries, it was necessary to generate ∼1,000,000 independent transformants to achieve ∼500,000 different Inr sequence variants because there are only 1,048,576 (= 4^10^) possible sequences in a randomized 10-mer, and we obtained many duplicate Inr sequences. The HARPE transcription reactions were scaled according to the number of independent transformants. To enable the use of HARPE for the Inr region, which includes sequence that are both upstream and downstream of the transcription start site, a downstream barcode was added by randomizing the region from +53 to +69 or +53 to +67 (relative to the SCP1 +1 TSS) for the 14-nt or 10-nt randomized Inr libraries, respectively, as was done for the HARPE analysis of the TATA box (Vo ngoc et al. 2020).

#### Transcription of HARPE libraries

In the biochemical HARPE experiments, the plasmid libraries were subjected to in vitro transcription with HeLa nuclear extracts (6 or 12 standard reactions for the 14-nt or 10-nt randomized Inr libraries, respectively; as noted above, the reactions were scaled according to the number of independent transformants in each library) as in Vo ngoc et al. (2020). The RNA from the 6 or 12 reactions was combined and processed as described below. In the cell-based HARPE experiments, the plasmid libraries were transfected into HeLa cells [five 10-cm culture dishes per library (14-nt randomized Inr libraries only)] as in Vo ngoc et al. (2020). During collection, one-fifth of each cell pellet was reserved for plasmid DNA extraction, and the remainder of the cells was used for RNA extraction. All HARPE experiments were performed independently at least two times to ensure reproducibility of the data. Replicates were performed with the same HARPE DNA libraries in independent transcription reactions or transfection analyses.

#### HARPE RNA extraction and processing

RNA transcripts from cells or from in vitro transcription reactions were extracted with Trizol or Trizol LS (Thermo Fisher Scientific), respectively. Total RNA (120 μg per condition for cell transfection experiments, or the entirety of each condition for in vitro transcription experiments) was processed as described in Vo ngoc et al. (2020, 2023). Briefly, the following steps were performed. First, contaminating plasmid DNA was removed with the TURBO DNA-free Kit (Thermo Fisher Scientific). Next, the RNA was subjected to reverse transcription with SuperScript III Reverse Transcriptase (Thermo Fisher Scientific) and treatment with RNase H (New England Biolabs). The resulting cDNA was size-selected on a 6% (w/v) polyacrylamide-8 M urea gel with fluorescently-labeled size markers. The size-selected cDNAs and HARPE plasmid DNAs were used as templates to generate DNA amplicons for Illumina sequencing (RNA and DNA datasets, respectively). In the cell-based HARPE experiments, the DNA datasets were obtained with plasmid DNA samples that were extracted post-transfection.

Custom forward oligonucleotides containing the Illumina P5 and Read1-primer sequences preceding the sequences corresponding to nucleotides +19 to +34 (relative to the SCP1 +1 TSS; the SCP1 DPR and mutDPR promoter backbones required different primers) were used to amplify the randomized barcode regions. Custom forward oligonucleotides containing the Illumina P5 and Read1-primer sequences preceding the sequences corresponding to nucleotides −44 to −23 or −40 to −23 (relative to the SCP1 +1 TSS; the SCP1 TATA and mutTATA promoter backbones required different primers) were used to pair each barcode with its corresponding Inr variant. Reverse primers were selected from the NEBNext® Multiplex Oligos for Illumina® kits (New England Biolabs), which match the Illumina Read2-primer sequence present on the HARPE plasmid and corresponding cDNA. NGS PCR amplicons were size-selected on native 6% (w/v) polyacrylamide gels prior to Illumina sequencing.

#### HARPE sequencing and data processing

Illumina sequencing of NGS PCR amplicons was performed on NovaSeq 6000 or NovaSeq X Plus sequencing systems at the IGM Genomics Center, University of California, San Diego (supported by NIH SIG grant #S10 OD026929). NGS data processing was performed as described in Vo ngoc et al. (2020, 2023). Paired-end (PE100) sequences were required to have a perfect match to the 10-nt sequences immediately upstream and downstream of the randomized Inr region, which was required to be exactly 10 nt or 14 nt (for experiments with the 10-nt or 14-nt randomized Inr libraries, respectively). Sequences were also required to have a perfect match to the 10-nt sequences immediately upstream and downstream of the randomized barcode region, which was required to be exactly 15 nt or 17 nt (for experiments with the 10-nt or 14-nt randomized Inr libraries, respectively). All reads containing a match to the selection pattern were deemed usable and trimmed for sequences outside the randomized region.

Conversion tables from barcode to Inr variant were built by paired-end sequencing of amplicons from the starting plasmid libraries. Both read 1 and read 2 were required to pass the above screening criteria. Barcodes that were associated with multiple Inr variants were discarded (this was rare given the complexity of the 15-nt and 17-nt regions used). DNA and RNA datasets for all Inr HARPE experiments were matched to their corresponding barcode-to-Inr conversion tables. For the DNA datasets, we only used sequences with a minimum absolute read count of 10 and a minimum relative count of 0.2 or 0.75 reads per million (RPM; for experiments with the 10-nt or 14-nt randomized Inr libraries, respectively) so that low-confidence variants would not be included in the analyses. RNA dataset sequences were then matched to their corresponding DNA dataset. The transcription strength for each Inr sequence variant was defined as the RNA read count (in RPMs) divided by the DNA read count (in RPMs). Inr variants associated with multiple barcodes were combined, and their transcription strengths were computed as the average transcription strength across multiple barcodes.

#### SVR model training and optimization

Machine learning analyses were performed as described in Vo ngoc et al. (2020, 2023), except that we used functions from the Python package scikit-learn (version 1.3.0; Pedregosa et al. 2011) instead of the R package e1071 (version 1.7-2). For SVR training, we used the default radial basis function (RBF) kernel, which yielded the best results among those tested.

Nucleotide variables for HARPE variants were computed as four categories (A, C, G, T) using one-hot encoding. These categories were used as the input features, and transcription strength was used as the output variable. For each SVR model, we set aside approximately 7,000-8,000 independent test sequences that represent the full range of transcription strengths. With the remaining sequences, we trained the SVR models with the 100,000 most active (best) variants and 100,000 randomly selected non-best variants. Grid search was performed for the hyperparameters cost and gamma, and cross validation was carried out with two independent test sets (two halves of the test sequences) that were not used for training (Supplemental Fig. S3). The selected hyperparameters for each model are as follows: for the Inr10 (with DPR) SVRb, Inr10 (with TATA) SVRb, Inr14 (with DPR) SVRb, and Inr14 (Inr only) SVRb models, we used cost = 10 and gamma = 0.1; for the Inr14 (with TATA) SVRb, Inr14 (with DPR) SVRc, and Inr14 (with TATA) SVRc models, we used cost = 1 and gamma = 0.1.

#### Motif discovery

Motif discovery was performed with Hypergeometric Optimization of Motif EnRichment (HOMER; Heinz et al. 2010). findMotifs.pl was used to search for de novo motifs among the most active (best) HARPE sequence variants. Randomly selected non-best sequence variants were used as background. WebLogo 3 (Crooks et al. 2004) was used to generate all sequence logos.

#### Identification of transcription start sites

Human transcription start sites (TSSs) were identified by using the GRO-cap method in GM12878 cells (Core et al. 2014; GSE60456) and csRNA-seq method in K562 cells (Duttke et al. 2019; GSE135498). *Drosophila* TSSs were identified by using the csRNA-seq method in *Drosophila* embryos (Delos Santos et al. 2022; GSE203135) and 5’-GRO-seq method in S2 cells (GSE68677). Focused TSSs in the human and *Drosophila* TSS data were identified as described in Vo ngoc et al. (2017) by using Focus_TSS.py. The corresponding promoter sequences were extracted from the human genome [hg19 (GM12878) and hg38 (K562)] and *Drosophila* genome [dm6 (embryos) and dm3 (S2)] by using homerTools extract (Heinz et al. 2010).

#### Gene ontology analyses

Human transcription start sites (TSSs) from the GM12878 GRO-cap dataset (Core et al. 2014) were annotated by using annotatePeaks.pl from the HOMER suite (Heinz et al. 2010) with the hg19 genome assembly and GENCODE Release 19 annotations. GRO-cap TSSs located >200 nt from the nearest annotated TSS were excluded. The genes for which the GRO-cap TSSs are ≤200 nt from the nearest annotated TSSs were used as the “universe” (background) gene list. Gene ontology (GO) enrichment analyses were performed by using the R package clusterProfiler (Wu et al. 2021), with gene-to-GO term mappings provided by the org.Hs.eg.db annotation package. Ensembl gene IDs and the Biological Process (BP) ontology were used. Over-representation analysis (ORA) was used to identify GO terms enriched among genes whose corresponding promoters contained either a TATA box only (TATA+, Inr−, DPR−) or an Inr and DPR (TATA−, Inr+, DPR+). The activity of each element was evaluated by using the SVR models of the human TATA box (Vo ngoc et al. 2020; GSE139635), human Inr [Inr10 (with DPR) SVRb model; this study], and human DPR (Vo ngoc et al. 2020; GSE139635). Approximately 4% of the “universe” genes contained a TATA box only (TATA+, Inr−, DPR−), and approximately 13% had an Inr and DPR (TATA−, Inr+, DPR+). ORA yielded 29 significant (adjusted *p* < 0.05) GO terms for TATA-only promoters and 0 (zero) significant GO terms for Inr+DPR promoters. Gene set enrichment analysis (GSEA) was also performed to identify GO terms with a skewed distribution across the full gene list ranked by TATA SVR score. This analysis identified 170 significant (adjusted *p* < 0.05) GO terms. The top 25 GO terms from the ORA and GSEA analyses are shown in Supplemental Table 4A and 4B, respectively.

Human TSSs from the K562 csRNA-seq dataset (Duttke et al. 2019) were annotated by using annotatePeaks.pl from the HOMER suite (Heinz et al. 2010) with the hg38 genome assembly and GENCODE Release 49 annotations. csRNA-seq TSSs located >200 nt from the nearest annotated TSS were excluded. For promoters containing a perfect match to the TCT motif (YYC_+1_TTTY), the associated genes are listed in Supplemental Tables 3A and 3B, ranked by Inr (with DPR) and Inr (with TATA) SVRc scores, respectively.

#### Data analysis, statistics, and graphical displays

All general data processing and visualizations were performed in R (version 4.4.0) with R packages tidyr (v1.3.1), dplyr (v1.1.4), ggplot2 (v3.5.1), stringr (v1.5.1), Biostrings (v2.72.1), or in Python (version 3.12.1) with Python packages numpy (v1.26.4), pandas (v2.2.3), scipy (v1.12.0), matplotlib (v3.10.0), seaborn (v0.13.2), and biopython (v1.83).

## Notes

### Competing Interest Statement

The authors have declared no competing interest.

### Summary of Updates

The revised manuscript contains new data (Supplemental Figure S21), revised figures (Figures 2C, 2D; Supplemental Figures S12 and S13), and updated text and conclusions, especially with regard to the TCT motif and its relationship to other core promoter elements in humans and Drosophila. In addition, we further highlighted the quantitative genome-wide nature of our analyses of natural human and Drosophila core promoters by using the SVR models.

